# Single cell analysis of the aging female hypothalamus

**DOI:** 10.1101/2021.03.07.434282

**Authors:** Kaitlyn H. Hajdarovic, Doudou Yu, Lexi-Amber Hassell, Shane Evans, Sarah Packer, Nicola Neretti, Ashley E. Webb

## Abstract

Alterations in metabolism, sleep patterns, body composition, and hormone status are all key features of aging. The hypothalamus is a well-conserved brain region that controls these homeostatic and survival-related behaviors. Despite the importance of this brain region in healthy aging, little is known about the intrinsic features of hypothalamic aging. Here, we utilize single nuclei RNA-sequencing to assess the transcriptomes of 40,064 hypothalamic nuclei from young and aged female mice. We identify cell type-specific signatures of aging in neurons, astrocytes, and microglia, as well as among the diverse collection of neuronal subtypes in this region. We uncover key changes in cell types critical for metabolic regulation and body composition, as well as in an area of the hypothalamus linked to cognition. In addition, our analysis reveals an unexpected female-specific feature of hypothalamic aging. Specifically, we discover that the master regulator of X-inactivation, *Xist*, is elevated with age, particularly in hypothalamic neurons. Moreover, using machine learning, we show that levels of X-chromosome genes, and *Xist* itself, are the best predictors of cellular age. Together, this study identifies critical cell-specific changes of the aging hypothalamus in mammals, and uncovers a novel marker of neuronal aging in females.

## INTRODUCTION

While human lifespan has increased dramatically in recent years, improvements in healthspan, the period of life in which a person is disease-free, have been more modest^1^. Susceptibility to a host of diseases increases with aging, including diabetes, stroke^2^, cancer^3^, and neurodegenerative diseases^4^. Aging is accompanied by changes in body composition, including decreased lean muscle mass, loss of bone density, and increased abdominal fat^1^. Concomitant with these changes are alterations in endocrine states, such as decreased sex hormone production, and reduced growth hormone and insulin-like growth factor-I^5^. Endocrine function and homeostatic processes, such as energy homeostasis^6^ and release of sex hormones^5^, are controlled by neuropeptidergic neurons in the hypothalamus.

Nutrient sensing is one of several functions of the hypothalamus that implicates this brain region in healthy aging. Specific neuronal subtypes in the hypothalamus respond to circulating cues to organize the response to dietary changes through regulation of energy balance, glucose homeostasis and growth factor secretion^6^. Caloric restriction (CR) is one of the most well-established behavioral interventions that improves lifespan and healthspan in many model organisms^7^. Genetic models that mimic the effects of CR via modulation of energy sensing pathways have revealed the mechanistic underpinnings of lifespan extension. For example, in *C. elegans*, the effects of dietary restriction are dependent on the function of neuropeptidergic energy sensing neurons; genetic manipulation of energy sensing genes in those neurons is sufficient to increase longevity^8^. Similarly, lifespan extension in the fruit fly *Drosophila* is dependent on specialized neurons called median neurosecretory cells^9^. In rodents, manipulations to the hypothalamus can also alter lifespan. Specifically, brain-specific over expression of *Sirt1* leads to alterations in the dorsomedial and lateral hypothalamus and increases lifespan^10^. In addition, alteration of immune signaling in the mediobasal hypothalamus affects longevity, with a reduction in immune signaling promoting longevity^11^.

Sex differences in lifespan have been documented in many species, including mice^12^. In humans, there is a robust difference in female and male lifespan across countries, with females living an average of 4-10 years longer than males^13^. Intriguingly, interventions that extend life span in model organisms do so in a sex-specific manner. For example, caloric restriction (CR) is one of the most robustly studied interventions and its effects have been observed from yeast to non-human primates^7^ Like many interventions, CR has sex-specific effects, with restricted females generally living longer than males on the same diet^14^. Similarly, the brain-specific *Sirt1* overexpression model results in a larger lifespan increase for females when compared to males^15^. However, the aging female brain remains critically understudied, and we know little about how areas involved in healthy aging, such as the hypothalamus, change with age in females.

Epigenetic and transcriptional changes are widespread across tissues during aging, including in the brain^16, 17^. Key transcriptional factors such as FOXO/DAF-16, NF-kB, and MYC function as conserved regulators of these networks and have been implicated in aging^11, 16, 18^. However, despite a great interest in how changes in transcriptional programs affect aging, our understanding of how distinct cellular subtypes change transcriptionally with age remains limited. Investigation of how transcriptional programs change in a cell-type specific manner in the hypothalamus will provide important insight into the aging process across tissues. Recent advances in single-cell RNA-sequencing (RNA-seq) have expanded our understanding of the diverse cell types that comprise the hypothalamus^19–26^. This approach allows the investigation of previously unappreciated transcriptional and functional diversity of this brain region. Here, we use a single nuclei RNA-seq approach to identify aging-associated transcriptional changes across the mouse hypothalamus, thereby capturing the diversity of cell types in this brain region.

## RESULTS

### Single nuclei sequencing of the aging mouse hypothalamus

We employed single nuclei RNA sequencing (snRNA-seq), which is currently the optimal method for single cell transcriptomic profiling of the diversity of cell types in the adult mammalian brain^27, 28^. We isolated nuclei from the hypothalamus of young (3 month) and aged (19-24 month) female mice, with four replicate libraries for each age (Figure 1A). After quality filtering, we obtained 40,064 high quality nuclei for analysis: 16,256 and 23,808 nuclei from young and aged animals, respectively (Figure S1A). Cellular composition and data quality were similar across replicates (Figure S1B-D).

**Figure 1.**
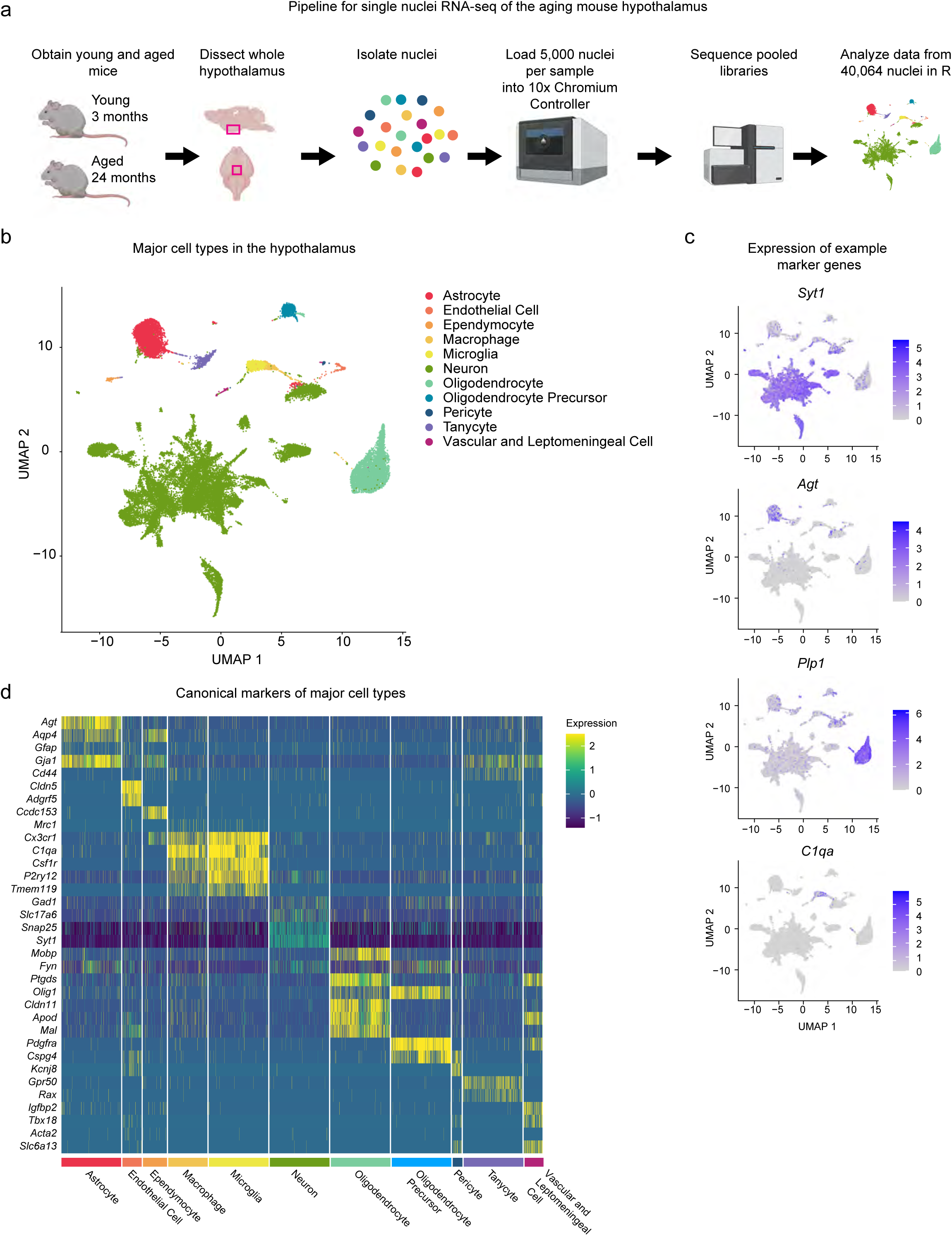
Single-nuclei analysis of the hypothalamus. A) Schematic detailing the experimental workflow from dissection through analysis. B) Uniform Manifold Approximation and Projection (UMAP) plot of all 40,064 nuclei used for analysis. Clustering analysis revealed 11 broad categories of cell type identity. C) UMAP plots of all nuclei labeled for expression of cell type-specific markers. *Syt1*, neurons; *Agt*, astrocytes; *Plp1*, oligodendrocytes; *C1qa*, microglia/macrophages. Color scale indicates level of gene expression. D) Heatmap highlighting expression of cell type markers in each cluster, a maximum of 500 nuclei per cluster are displayed.

Clustering analysis with the Louvain algorithm revealed distinct clusters representing the major cell types of the hypothalamus, which we identified based on expression of canonical markers (Figure 1B-D). For each individual cluster, we identified the top 10 genes that were differentially expressed using the Wilcoxon Rank Sum test (Supplementary Table 1). For example, neurons were defined by expression of *Syt1*, astrocytes defined by *Agt and Gja1*, oligodendrocytes by *Olig1* and *Plp1* expression, and oligodendrocyte precursor cells (OPCs) were identified by expression of *Pdgfra*. The microglia and macrophage clusters were defined by expression of *C1qa* and distinguished by higher expression of *Tmem119* and *P2ry12* in the microglia cluster (Figure 1C). Less abundant cell types were also observed, including ependymocytes (ependymal cells; *Ccdc153*), pericytes (*Flt1*), endothelial cells (*Clnd5*), and vascular and leptomeningeal cells (VLMC; *Slc6a13*). We also observed a distinct cluster of tanycytes, which are specific to the hypothalamus and defined by *Rax* expression. Nuclei in these broad categories expressed additional canonical markers associated with their cell type, for example, the astrocyte cluster expressed *Gfap,* further validating the identify of each cluster.

Cell type diversity is achieved through expression of transcriptional regulators that orchestrate cell type-specific gene expression networks. To identify the regulators responsible for distinct expression networks across cell types in the hypothalamus, we used SCENIC, a regulatory network inference tool^29^. SCENIC identifies regulons, defined as a transcription factor and the genes it regulates, and scores the activity of the regulons in individual cells. Further, it provides a regulon specificity score, which indicates whether a given regulon is specific to an individual cell type or shared among clusters. In our analysis, we observed strong shared and cell-type specific signatures for each cluster. For example, the Dbx2 regulon is strongly enriched in the astrocyte cluster (RSS = 0.348), with almost all astrocytes expressing the regulon, whereas, the Atf2 and Creb3l1 regulons are enriched in neurons (Figure Supplementary 2A-B, Supplementary Table 2) (RSS = 0.639 and 0.626, respectively). Together, this analysis identifies the distinct gene expression signatures in the major cell types in the hypothalamus that are orchestrated by specific combinations of transcriptional regulators associated with cell type identity.

### Major cell types of the hypothalamus acquire cell type-specific gene expression changes with age

We next investigated the changes in gene expression that occur with age in the major cell types of the hypothalamus. As expected, aging was not associated with changes in composition of this brain region, and each major cell type was similarly represented in young and aged mice (Figure S1C-D). To gain a global understanding of how gene expression is altered with age, we first performed differential expression analysis on all cells using the Model-based Analysis of Single-cell Transcriptomics (MAST)^30, 31^, with a random effect for sample of origin. Using this approach, we identified 275 and 342 genes that were upregulated and downregulated with age, respectively (padj < 0.05, fold change > 0.1) (Figure 2A, Supplementary Table 3). As an initial validation, we cross-checked our results with publicly available bulk microarray data on the aging hypothalmus^32^. Although these data differ from ours in regard to strain and sex, this analysis confirmed several changes in our dataset, including downregulation of *Gria1*, *Apoe*, *Camk2a*, and *Atp1b2* (Supplementary Table 3). Moreover, we confirmed a key finding from work on male rats showing that oxytocin binding is decreased in several brain regions with age, including the hypothalamus^33^. Here, we show a reduction in *Oxt* expression in aged female mice, suggesting a reduction of oxytocin signaling is a feature of both male and female aging. Interestingly, the most upregulated genes in the MAST analysis included *Xist* and *Tsix*, which are both long non-coding RNAs involved in X chromosome inactivation^34, 35^, and are female specific.

**Figure 2.**
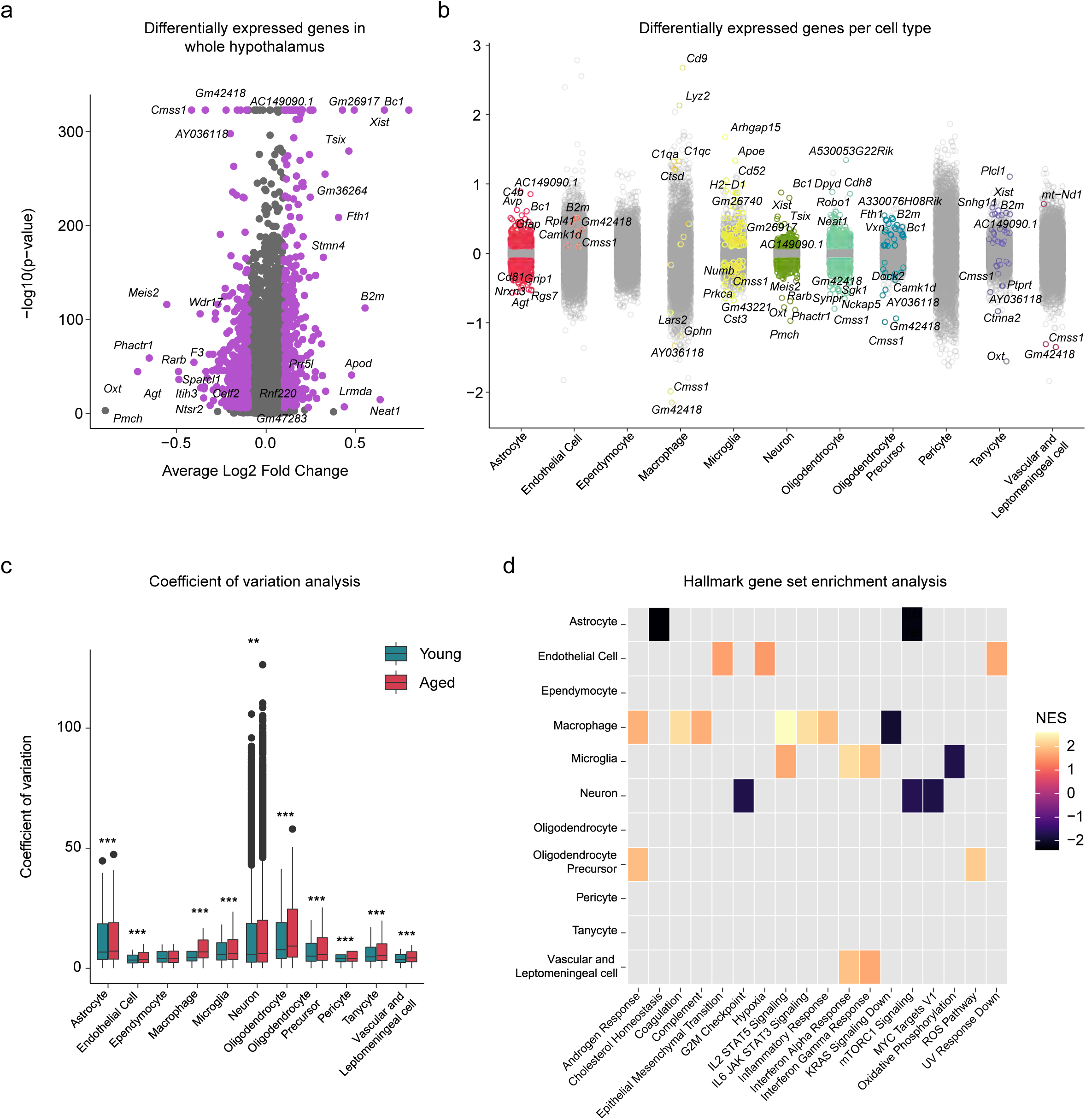
The aging hypothalamus harbors cell type-specific transcriptional changes. A) Volcano plot showing overall differential expression of genes between all young and aged nuclei. Significant genes in purple (adjusted p value < 0.05, FC > 0.1). Genes with p value of zero given arbitrarily small p value for plotting purposes. B) Strip plot showing DE genes in each cell type. Significant genes (adjusted p value < 0.05, FC > 0.1, MAST analysis with random effect for sequencing depth and sample of origin) in color, nonsignificant genes are in gray. Top 5 upregulated and top 5 downregulated genes per cluster labeled. C) Coefficient of variation analysis for each cellular subtype. In almost all subtypes the CV is significantly higher in the aged condition (Wilcoxen Test with Bonferonni correction, *** padj < 0.001, ** padj < 0.01 D) Heatmap showing GSEA enrichment analysis for Hallmark terms. Color indicates normalized enrichment score. Significant gene sets calculated as adjusted p value < 0.1.

Next, we investigated the impact of age on gene expression in each major cell type. Neurons, astrocytes, oligodendrocytes, and microglia showed the greatest numbers of differentially expressed genes with age (Figure 2B, Supplementary Table 4). Additionally, we performed coefficient of variation analysis on the major cell types and observed a significant difference between ages, with nearly all types showing in increase with age (Figure 2C). This finding suggests that variability in gene expression increases with age in most cell types, which likely contributes to cellular dysfunction within the aged hypothalamus.

To validate our findings and determine the extent to which the changes we observe are specific to the hypothalamus, we compared our dataset to a publicly available snRNA-seq dataset analyzing the female mouse hippocampus^36^. Astrocytes, oligodendrocytes, and microglia all showed high agreement between the hypothalamic and hippocampal datasets, for example, both datasets show significant upregulation of *C4b* in astrocytes, *Apoe* and *Lyz2* in microglia, and *Cdh8* and *Neat1* in oligodendrocytes (Figure 3A). When we compared the relationship in log2 fold change of significant (padj < 0.05) genes between the hypothalamus and hippocampus, a statistically significant positive correlation emerges for astrocytes (ρ = 0.78, p < 0.001), oligodendrocytes (ρ = 0.61, p < 0.001), and microglia (ρ = 0.85, p < 0.001). Interestingly, in contrast to glia, there was less overlap in differentially expressed genes between the hypothalamic neurons and hippocampal neurons, and there was little correlation in gene expression changes between hypothalamic neurons and hippocampal (ρ = -0.07, p < 0.001). While some genes, such as *Xist,* are upregulated with age in both neuronal subsets, other genes are either unchanged or regulated in the opposing direction in the two sets (Figure 3B). For example, *Phactr1* and *Meis2* are among the most significant downregulated neuronal genes in the hypothalamic dataset but are not significantly changed with age in the hippocampal dataset (padj > 0.05). The genes *Rsrp1*, *Gm26917*, and *Kcnip4* are upregulated with age in the hypothalamus, but downregulated in the hippocampus with age. Intriguingly, the gene *Rps29* is upregulated in both datasets, in agreement with previously single cell RNA sequencing studies of the aged mouse brain^37^. Together these data suggest that glia share harbor general signatures of aging in distinct brain regions, whereas and region-specific signatures are predominant in neuronal aging.

**Figure 3.**
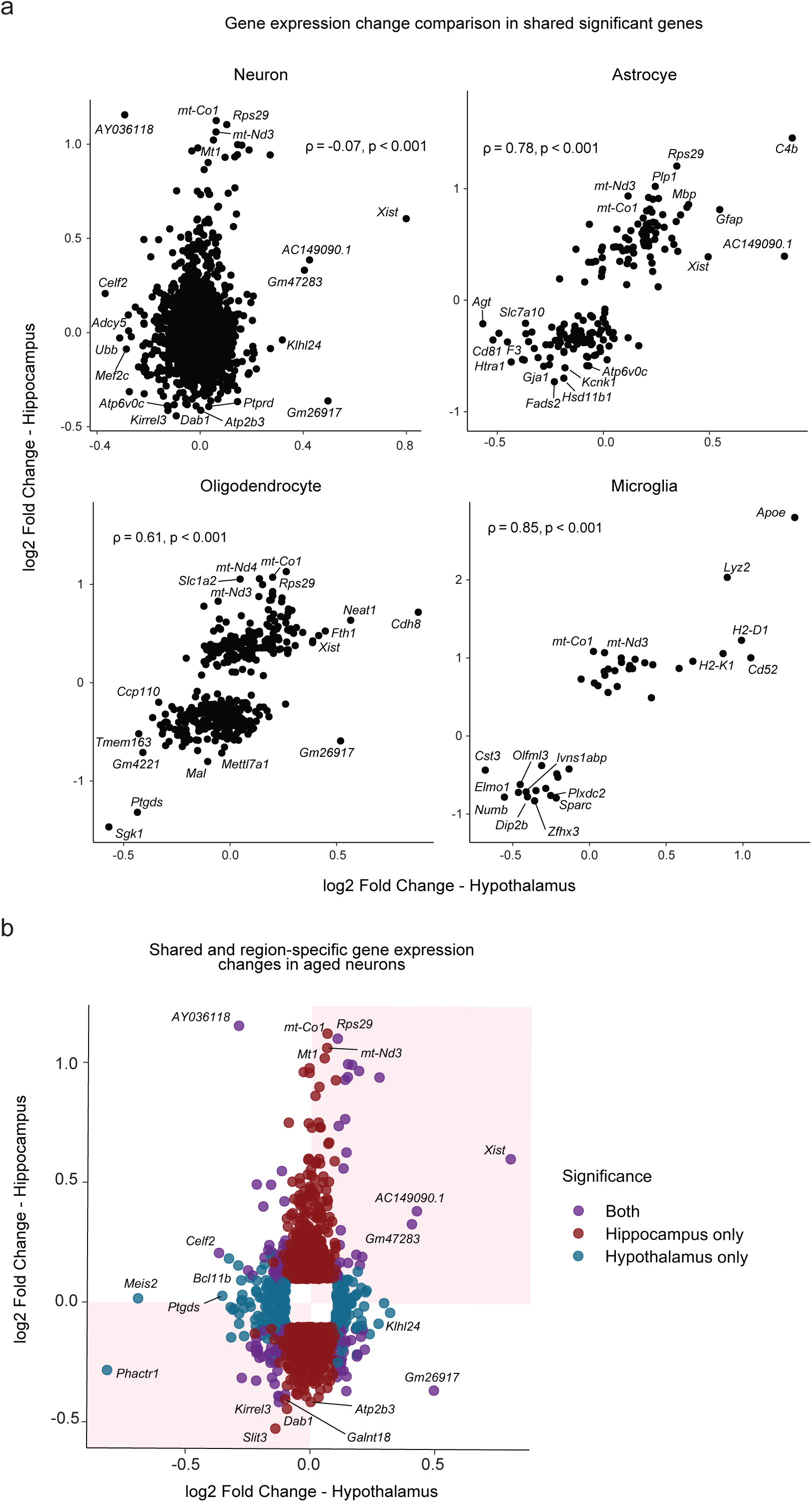
Shared and region-specific aging signatures between hypothalamus and hippocampus. A) Spearman’s rank correlation of log2 fold change in significant (padj < 0.05) genes shared between the hypothalamus and hippocampus. Correlation of expression change in neurons is very weakly negatively correlated (ρ -0.069, p <0.001) while gene expression changes in other cell types are strongly positively correlated: astrocyte (ρ = 0.78, p < 0.001), oligodendrocyte (ρ = 0.61, p < 0.001), microglia (ρ = 0.85, p < 0.001). B) Scatterplot of genes significant in at least one dataset (padj <0.05, log2 fold change > 0.1). Color indicates whether a gene is significant in both datasets (purple), hypothalamus only (blue) or hippocampus only (red). Shading indicates genes that are changing in the same direction for both datasets.

To investigate the cellular processes that are altered with age in the different cell types in the hypothalamus, we performed Gene Set Enrichment Analysis (GSEA) using the hallmark gene set^38^ (Figure 2D). We observed unique signatures of aging in each cell type, as well as some shared hallmarks of aging. For example, neurons and astrocytes share an under-enrichment (negative normalized enrichment score) for the mTORC1 signaling gene set. Astrocytes were also under-enriched in genes related to cholesterol homeostasis, which aligns with previous research showing a decrease in expression of cholesterol synthesis genes in aged hypothalamic astrocytes^39^. In contrast, neurons displayed alterations in G_2_M checkpoint genes, which is particularly interesting in light of evidence linking Alzheimer’s Disease to aberrant cell cycle entry driven through activation of mTORC1^40^. Interestingly, mTORC1 activation in the arcuate nucleus of the hypothalamus is reduced in ovariectomized mice^41^. Thus, mTORC1 under-enrichment in the aged female hypothalamus represents a key a target for further study into the intersection of sex, aging, and neurodegenerative disease.

Microglia and macrophages function as immune cells in the brain, and both cell types show striking cell type-specific gene set enrichments with age (Figure 2D). Macrophages are enriched in IL2 STAT5 signaling, IL6 JAK STAT3 signaling, and inflammatory response gene sets. Microglia are enriched in genes related to the interferon alpha response and interferon gamma response, and are under enriched in the oxidative phosphorylation gene set with age. These data suggest distinct changes in different subsets of immune cells in the aging hypothalamus.

Glial cells, including microglia are critical regulators of neuronal function. To understand how the relationships between these cells and neurons are changed with age, we utilized the ligand-receptor repository CellPhoneDB^42^ to infer cell-cell interactions (Supplementary Figure 3A). Interestingly, we found that loss of ligand-receptor interactions involving growth factors was a theme across the cell types studied. For example, in the young astrocyte-neuronal and tanycyte-neuronal cell pairs, TGFB2_TGFBR3 was enriched. Both cell pairs lose this enrichment with age. Similarly, there is a loss of enrichment in pathways involving FGF9. Specifically, while FGF9-FGFR3 and FGF9-FGFR1 are enriched in young astrocyte-neuronal and tanycte-neuronal cell pairs, this enrichment is lost with age. The astrocyte-neuronal and oligodendrocyte-neuronal pairs also lose enrichment of the FGF9-FGFR2 ligand-receptor interaction with age. These factors are involved in diverse processes such as repair, learning and memory, and neurogenesis^43^. Additionally, the aged astrocyte-neuron cell pair loses SEMA3A_NRP1 signaling with age, while the microglia-neuronal cell pair loses both SEMA3A-NRP1 enrichment and SEMA3A-PlexinA4 enrichment. Semaphorin3a is a known player in synaptogenesis^44^. Taken together, the loss of these cell signaling pathways may represent a mechanism for alternations in synaptogenesis and neuronal homeostasis in the aged hypothalamus.

### Aged hypothalamic microglia are heterogeneous, representing a progressive aging trajectory

Microglia are macrophage-like cells found throughout the brain, and are critical for the immune response, including release of cytokines and chemokines, antigen presentation, and phagocytosis of debris^45^. Recent studies have revealed gene expression changes and microglial activation in the aged brain, which likely contribute to neurodegeneration^45^. Based on our findings that microglial-neuronal interactions involving the Alzheimer’s associated gene APP, as well as MIF were enriched in age (Supplementary Figure 3A), we sought additional strategies to uncover changes in these cells over time. Using Monocle3^46^, we performed pseudotemporal ordering of nuclei from the microglia and macrophage clusters. The trajectory accurately captures the transition from young to aged nuclei, suggesting a gradual progression toward aging in this cell type, and a significant increase in the proportion of aged nuclei across pseudotime (Figure 4A-B). To confirm this analysis, we freshly isolated CD11b+ hypothalamic microglia from mice at three timepoints (3, 12, and 24 months) and performed qPCR for candidate genes discovered in the pseudotime analysis. This experiment recapitulated specific genes trajectories (Supplementary Figure 4A), validating this approach.

**Figure 4.**
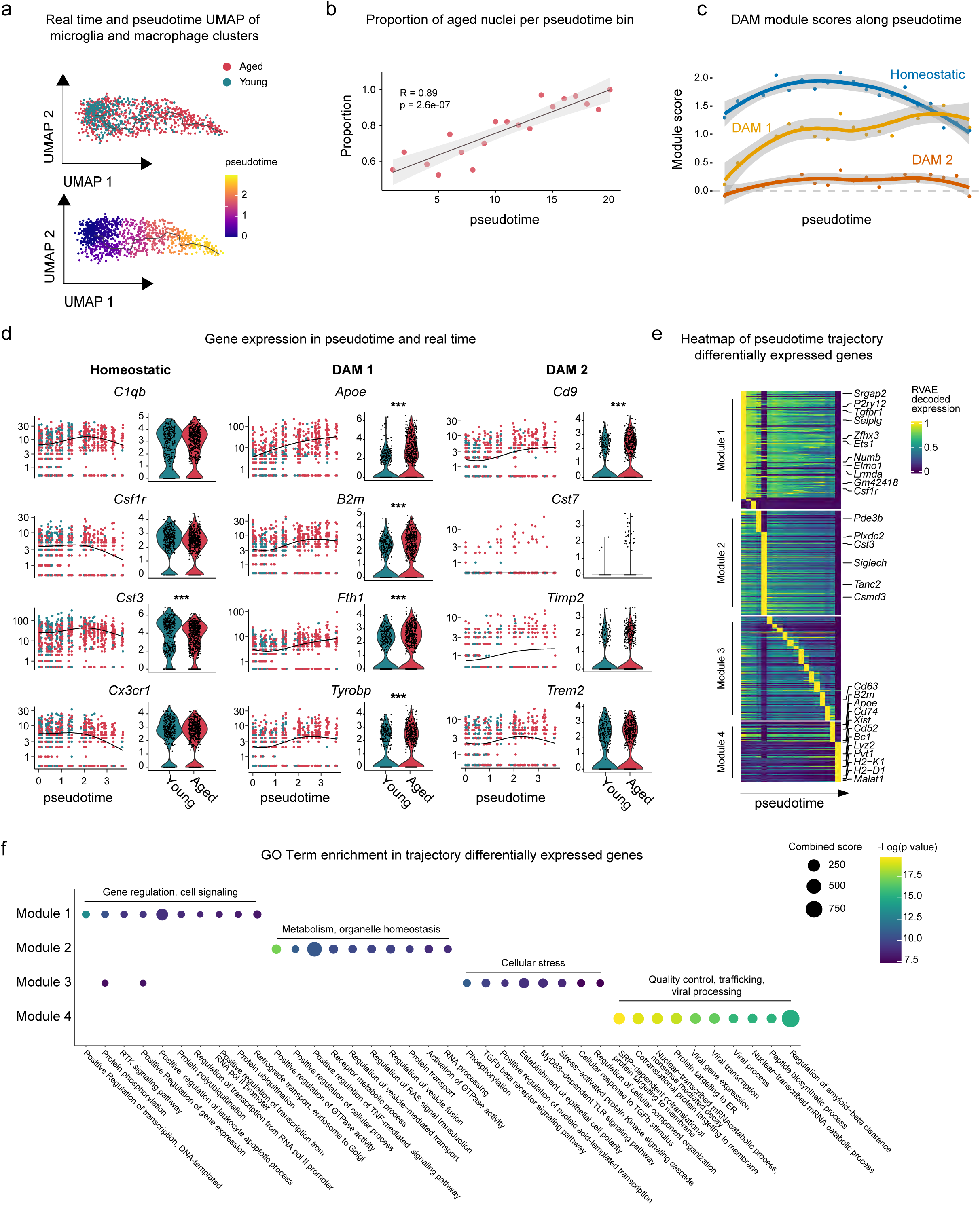
Trajectory analysis of aging hypothalamic microglia. A) Monocle3 pseudotemporal ordering of microglia and macrophage clusters (n = 1121 nuclei) defining a single trajectory from young to aged nuclei. Nuclei are colored by age (top) and pseudotime (bottom). B) Scatterplot showing the proportion of aged nuclei along the pseudotime timeline in 20 time bins (sized 0.15 per time bin). Pearson correlation of the proportion of aged nuclei and pseudotime timeline, R = 0.89 and p-value = 2.6e-7, 95% confidence interval shown. C) Plot showing the module expression score of three microglia states (homeostatic, DAM 1, and DAM 2). The darker lines are the local regression result for individual time bins (20 total), with the gray shadow depicting the 95% CIs. D) Left: kinetics plot showing the relative expression of representative genes for microglia states. The lines approximate expression along the trajectory using polynomial regressions. Right: violin plots of gene expression and MAST (*, adjusted p value < 0.05, ***, adjusted p value < 0.001). E) Heatmap showing modules of trajectory differentially expressed genes (t-DEGs) in the microglia cluster (n = 2112 genes). The expression value is RVAE decoded expression. The genes were grouped into four modules after ranking by the RVAE decoded expression. Module 1 (645 genes), module 2 (570 genes), module 3 (566 genes), and module 4 (331 genes). F) Dot plot showing the top 10 GO biological process terms for genes in individual modules.

Since changes in microglia have been implicated in both physiological aging and neurodegeneration, we examined how disease associated microglia genes (DAM genes) change as a function of pseudotime. We aggregated the expression of key genes from three microglia gene sets identified in the literature^47, 48^: homeostatic microglia (homeostatic), TREM2 independent stage 1 DAM (DAM 1), and TREM2 dependent stage 2 DAM (DAM 2), and plotted the aggregated expression as a module score along the pseudotime trajectory. Overall, there was a decrease in the module score for the homeostatic module over pseudotime suggesting a loss of maintenance of healthy microglia over time. In contrast, we observed an increase in the DAM 1 disease module score near the end of pseudotime (Figure 4C). The DAM 2 module does not seem to play a role in steady-state hypothalamic aging, as the module score remains low throughout pseudotime.

To further understand the role of these gene modules in aging hypothalamic microglia, we visualized gene expression across psuedotime and through real time (Figure 4D). While young microglia generally cluster earlier in pseudotime (pseudotime 0.0 through 1.5), aged microglia expressing these genes are distributed throughout pseudotime. Thus, hypothalamic microglia from aged animals have increased heterogeneity representing a progressive aging trajectory. While a subset of aged microglia retaining a youthful gene expression signature, many aged microglia highly express disease associated genes.

To fully capture gene expression changes along the trajectory, we performed Moran’s I test on microglia and macrophage genes, and found 2,112 statistically significant trajectory-dependent genes (Supplementary Table 5). To characterize their expression dynamics along pseudotime, we applied RVAgene^49^, an autoencoder neural network framework to reconstruct and smooth the pseudotime-dependent genes expression. We then visualized the RVAE decoded expression along pseudotime in a heatmap, and manually grouped the genes into four modules according to their pseudo-temporal expression patterns (Figure 4E). For example, genes in the module 1 are highly expressed in early pseudotime while genes in the module 4 are expressed in late pseudotime. To understand the biological processes enriched in each module, we performed GO analysis (Figure 4F). Interestingly, gene modules are transitioning through pseudotime from positive regulation of biological processes to immune responses, and finally to the amyloid-beta clearance and viral infection corresponding to known phenotypes of normal aging.

### Age-associated changes in X-inactivation genes is a sexually dimorphic feature of aging

Our initial differential expression analysis revealed the unexpected finding that the long non-coding RNA *Xist* is one of most highly upregulated genes in the female hypothalamus with age (Figure 3A). Differential expression analysis of each major cell type indicated upregulation of *Xist* with age in astrocytes, macrophages, microglia, neurons, oligodendrocytes, as well as tanycytes (Figure 5A), and we observed upregulation of *Xist* in the aging hippocampus as well^36^ (Figure 3A-B). *Xist* is a key player in X chromosome inactivation in females and is encoded on the X-inactivation center (XIC), which harbors additional non-coding RNA genes involved in the same process^34, 35, 50^. Intriguingly, we observed age-related upregulation of related RNAs in some cell types: *Ftx, Jpx,* and, *Tsix* (Figure 5A). We validated the upregulation of *Xist* using RNA extracted from independent tissue samples of different brain regions (hypothalamus, cerebellum, cortex and olfactory bulb). Strikingly, although *Xist* trended up in all brain regions we tested, the upregulation only reached significance in the female hypothalamus, revealing a novel feature of female hypothalamic aging (Figure 5B). As expected, we did not detect *Xist* expression in adult male mice, and there was no upregulation of this gene with age in males (Figure 5B). We further confirmed this finding using RNAScope to detect the *Xist* transcript *in situ* in coronal sections through the mouse hypothalamus. The average intensity of *Xist* expression in aged female hypothalamus (25 months) was significantly higher than in young female hypothalamus (3 months) using this method (Figure 5C).

**Figure 5.**
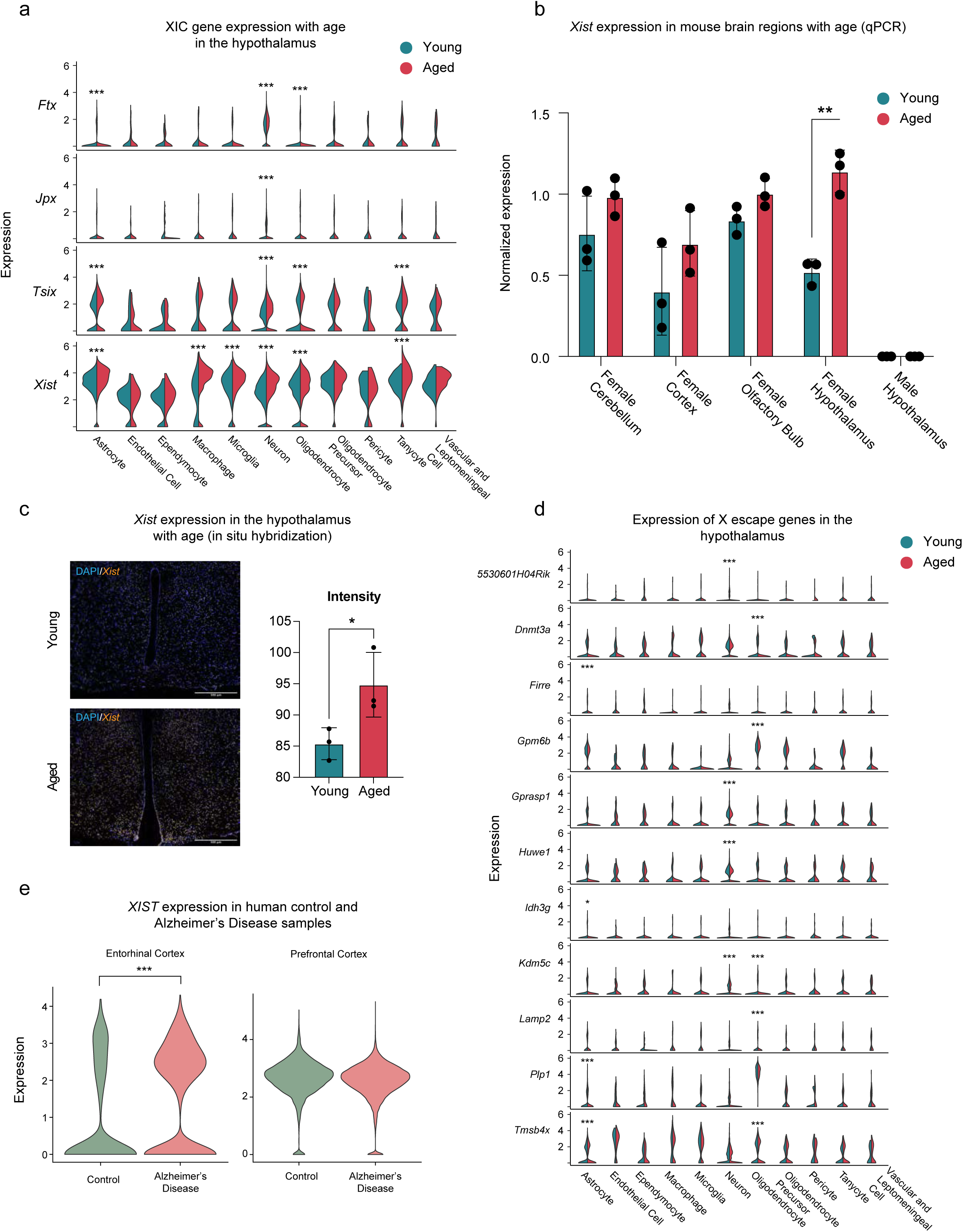
Alterations to X chromosome inactivation center are a feature of the aged female hypothalamus. A) Expression of genes involved in X chromosome inactivation by age and cell type. Differential expression between young and aged samples was calculated using MAST (*, adjusted p value < 0.05, ***, adjusted p value < 0.001). B) RT-qPCR of *Xist* expression in specific brain regions. *Xist* expression is significantly higher in the hypothalamus (n = 3 per age group, **p = 0.008, unpaired t-test with Bonferroni-Dunn correction). C) left, representative images from RNAScope for *Xist* transcript in young and aged female hypothalamus. Right, quantification of signal intensity (n = 3 animals per group, *p = 0.0473, unpaired t test). D) Expression of genes known to escape X chromosome inactivation by age and cell type. Differential expression between young and aged samples was calculated using MAST (*, adjusted p value < 0.05, ***, adjusted p value < 0.001). E) *XIST* expression in human entorhinal cortex (left) and prefrontal cortex (right). ***, adjusted p value < 0.001, MAST.

Although most genes on the inactive X chromosome are not expressed in females, a small number of genes are well-known to “escape” inactivation, and are expressed from both X chromosomes. These X escape genes are species and tissue specific^51, 52^. In the mouse, 14 genes escape X inactivation in brain tissue^51^.This list includes both *Xist*, and, *Ftx*, which have increased expression with age in our dataset. To determine if increased XIC gene expression with age might be affecting escape genes, we interrogated expression of genes known to escape X inactivation in mice. We compiled a list of genes that are both known to escape X inactivation in any tissue context in mice and are expressed in our dataset. We found that although changes to XIC genes seems to be uniform across cell types in our data, age-related changes to expression of X escape genes are cell type-specific (Figure 5D). For example, in astrocytes, *Idh3g* is downregulated with age, while *Firre*, *Plp1*, and *Tmsb4x* are upregulated. In neurons, *Gprasp1* and *Huwe1* are downregulated with age, while *5530601H04Rik* and *Kdm5c* are upregulated. Of the 39 X escape genes expressed in the dataset, 15 were differentially regulated with age in at least one cell type. These data suggest global changes to X chromosome regulation may be a feature of female hypothalamic aging.

Finally, to understand whether the changes in *Xist* we observed in mouse aging might be related to age-associated pathologies humans, we assessed changes in *XIST* expression between control and Alzheimer’s Disease human brain samples across two brain regions using publicly available snRNA-seq datasets^53, 54^. Using MAST with a random effect for sample of origin, we compared *XIST* expression across all cells from females in two independent datasets (Figure 5E). Strikingly, *XIST* is upregulated in human entorhinal cortex in women with Alzheimer’s, which is one of the earliest and most affected regions in this disease (log2 fold change = .574, padj < 0.001, n = 3942 nuclei). In contrast, nuclei derived from human prefrontal cortex shows no changes in *XIST* expression between control and Alzheimer’s Disease samples (padj > 0.05, n = 26212 nuclei). Thus, changes in *XIST* expression may be a brain-region specific feature of Alzheimer’s Disease in female patients.

### Neuronal subtype specific changes during aging

Hypothalamic neurons are highly diverse and function to orchestrate a wide range of processes and behaviors necessary for organismal survival^55^. This diversity of function is accomplished by cell type-specific gene expression programs, with each area of the hypothalamus containing a range of transcriptionally dissimilar neuronal subtypes^19–26^. Indeed, even neurons expressing the same neuropeptide gene may comprise functionally distinct subpopulations^6^. To address this complexity, we sub-clustered the neuronal nuclei to identify transcriptionally distinct populations. This analysis revealed 35 transcriptionally distinct clusters (Figure 6A), and broadly separated the nuclei into inhibitory (*Gad1* expressing GABAergic) or excitatory (*Slc17a6/*vGLUT2 expressing glutamatergic) identity (Figure 6B). The 35 clusters represent both known and undefined neuronal subtypes (see Supplementary Table 6 for markers of cluster identity). To discern the relationship between the clusters, we organized them according to transcriptional similarity using a Cluster Tree analysis (Figure 6C, left). Neurons with similar functions did cluster closely to one another. For example, some AgRP/NPY neurons and POMC neurons may arise from common progenitors^6^, and the Sst/Npy (29, expressing *Agrp*) and Pomc/Tac2 (31) clusters are near to one another on the cluster tree.

**Figure 6.**
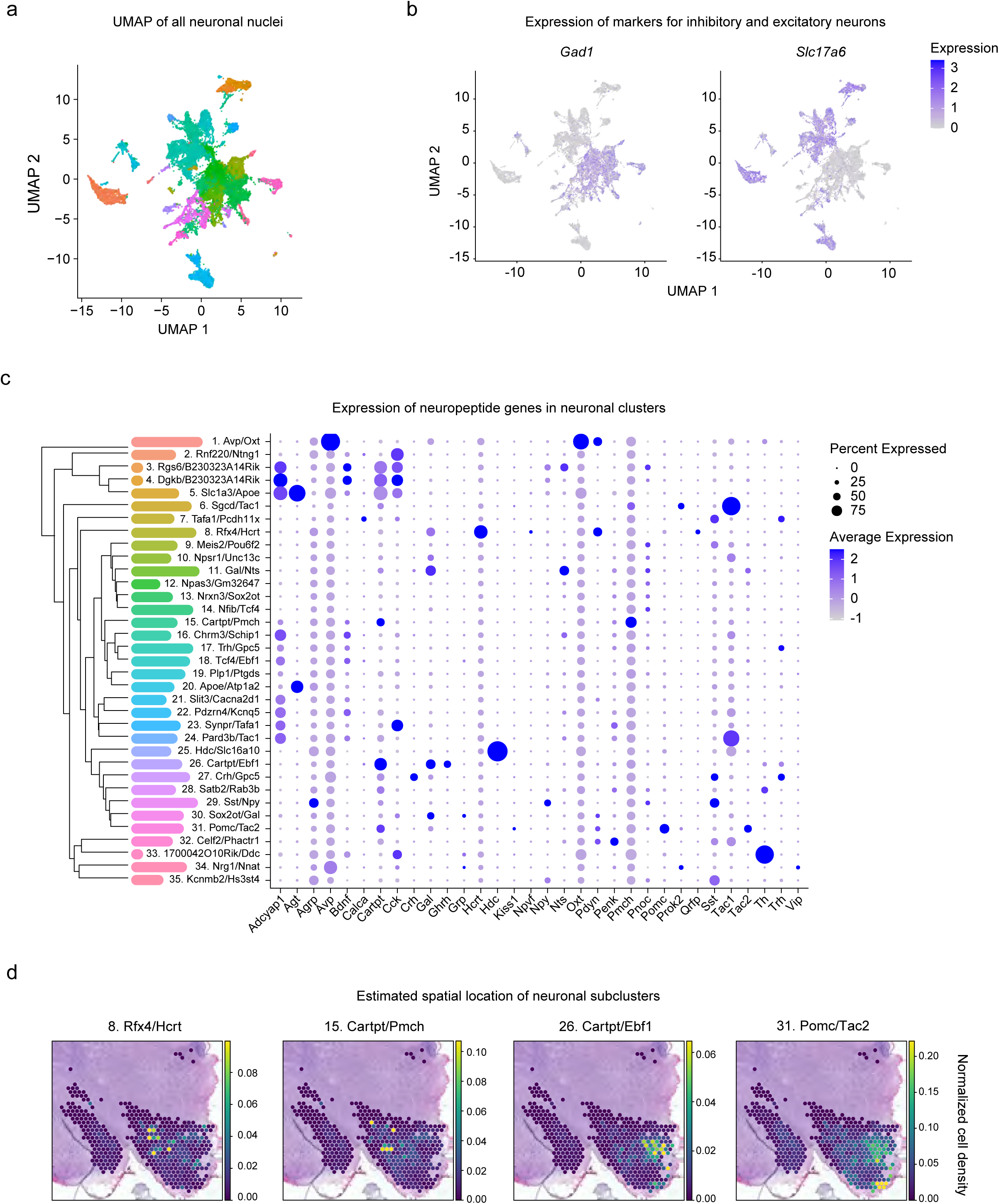
Identification of transcriptionally distinct neuronal subtypes. A) UMAP of all neuronal nuclei. Distinct clusters are identified by color, with identities listed in (C). B) UMAPs highlighting GABAergic (*Gad1*) and glutamatergic (*Slc17a6*) nuclei neuronal clusters. Color scale indicates expression level. C) left, Neuronal clusters are labeled according to the top 2 marker genes and ordered based on overall transcriptional similarity. Right, expression of neuropeptide genes in each cluster. Dot size indicates percent of nuclei expressing the gene, color indicates intensity of expression. D). Estimated spatial locations of neuronal subclusters.

We next investigated expression of specific neuropeptide genes across the clusters to functionally define the distinct neuronal subpopulations (Figure 6C, right). These clusters generally correspond to known cell types expressing one or two hallmark neuropeptides. We were able to identify neuronal clusters expressing genes encoding neuropeptides controlling processes which are altered with age (Supplementary Table 7). For example, we observed significant changes in clusters associated with feeding and energy homeostasis^56^, including those expressing the peptides agouti-related peptide (*Agrp*), Cocaine and amphetamine related transcript (*Cartpt*), Cholecystokinin (*Cck*), neuropeptide Y (*Npy*), proopiomelanocortin (*Pomc*), galanin (*Gal*), and hypocretin/orexin (*Hcrt*). Based on neuropeptide gene expression, these clusters most likely represent known neuronal populations with defined functions. For example, cluster Sst/Npy (29) is most likely comprised of AgRP/NPY neurons from the arcuate nucleus of the hypothalamus.

To further confirm neuronal subtype identity, we compared our dataset with publicly available spatial transcriptomic data from cell2location^57^. While mRNA signatures from broad cluster categories such as astrocytes (Supplementary Figure 5) do not show restriction to one or more hypothalamic subnuclei, the mRNA signatures of specific neuronal subclusters are localized in discrete locations. (Figure 6D, S4A-B). For example, the Pomc/Tac2 (31) cluster localizes to the most ventral portion of the coronal section (Figure 6D). Interestingly, two clusters expressing *Cartpt* (15. Cartpt/Pmch and 26. Cartpt/Ebf1) show little spatial overlap despite their shared neuropeptide profile, highlighting the strength of this method to define cell types both spatially and transcriptionally. Thus, this spatial analysis further validates the identity and function of the identified neuronal subclusters.

We next performed differential expression on clusters in which there were at least 20 nuclei per condition (Figure 7A, Supplementary Table 8). For each cluster, we also performed GSEA using the KEGG gene set. Most clusters tested exhibited significant transcriptional changes with age, although the number of DE genes varied by subtype. We observed that clusters expressing peptides involved in feeding and energy homeostasis were particularly altered with age in this analysis (such as 15. Cartpt/Pmch, 34 DE genes; 29. Sst/Npy, 65 DE genes; and 31. Pomc/Tac2, 42 DE genes). Among the many transcriptional changes found in the Pomc/Tac2 cluster (31) there was an intriguing downregulation of *Pcsk1n*. *Pcsk1n* encodes Proprotein Convertase Subtilisin/Kexin Type 1 Inhibitor, also called proSAAS, a propeptide which inhibits processing of other neuropeptides such as POMC^58^. This gene was also downregulated in a cluster of neurons expressing *Cartpt* (Cartpt/Ebf1 (26)). Interestingly, in a different *Cartpt* expressing cluster (Cartpt/Pmch (15)), the gene is upregulated with age, suggesting that changes to neuropeptide processing pathways with age are cell-type specific.

**Figure 7.**
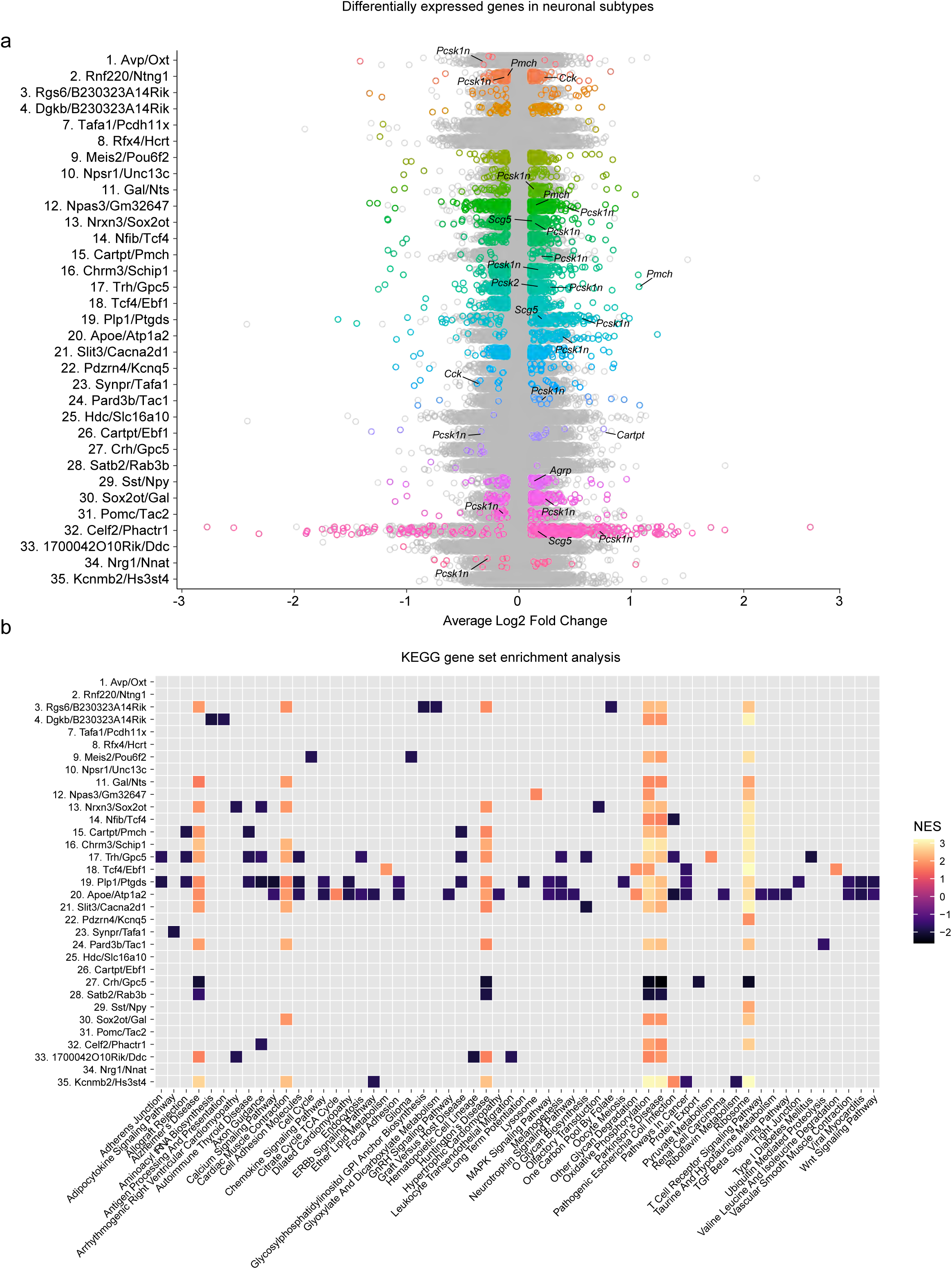
Neuronal subtypes exhibit distinct transcriptional changes with age. A) Strip plot showing DE genes per cluster. Significant genes (FC > 0.1, padj < 0.05) are colored, non-significant genes in gray. Genes discussed in text are labelled. B) Heatmap of GSEA results for each neuronal cluster. Significantly enriched terms (padj <0.1) are colored according to the normalized enrichment score.

Changes to expression of neuropeptide genes was also evident, with upregulation of *Agrp* in cluster Sst/Npy (29), upregulation of *Cartpt* in Cartpt/Ebf1 (26), and downregulation of *Cck* in two *Cck*+ subclusters (Synpr/Tafa1 (23) and Rnf220/Ntng1 (2)). Thus, for the first time, our dataset links neuron-specific gene expression changes in the hypothalamus with key features of organismal aging, such as weight and metabolic changes.

Based on expression of specific peptide genes (*Adcyap1*, *Cartpt*, *Cck*) and other established marker genes (*Foxb1*, *Cpne9*)^59^, we identified three clusters representing the medial mammillary nucleus of the hypothalamus: Rgs6/B230323A14Rik (3), Dgkb/B230323A14Rik (4), and Slc1a3/Apoe (5). This region is notable because unlike most areas of the hypothalamus, it is involved in memory via connections with the hippocampus^60^. While cluster Slc1a3/Apoe (5) had too few cells to meet our criteria for performing differential expression, both cluster Rgs6/B230323A14Rik (3) and Dgkb/B230323A14Rik (4) were significantly altered with age (Figure 7A). Gene set enrichment analysis using the KEGG gene set revealed enrichment for genes related to Alzheimer’s disease, cardiac muscle contraction, Huntington’s disease, oxidative phosphorylation, Parkinson’s disease, and the ribosome. There was an additional de-enrichment in genes related to glycosylphosphatidylinositol (GPI)-anchor biosynthesis and glyoxylate and dicarboxylate metabolism (Figure 7B). The identification of changes in this brain region is significant, as they may contribute to cognitive impairments with age.

Through our gene set enrichment analysis, a shared aging signature emerged among many hypothalamic neuronal subtypes. This included enrichment in pathways related to Alzheimer’s Disease (14 clusters), Huntington’s Disease (11 clusters), oxidative phosphorylation (19 clusters), Parkinson’s Disease (17 clusters), and the ribosome (21 clusters) (Figure 7B). A notable exception to this signature is the cluster most likely representing corticotropin releasing hormone (CRH) neurons of the paraventricular nucleus of the hypothalamus (Crh/Gpc5 (27)). CRH neurons are an integral component of the hypothalamic-pituitary-adrenal axis in the stress response^61^. Decreased CRH has been studied for several decades as a potential hallmark of Alzheimer’s Disease^62^, and CRH itself has been shown to be neuroprotective against Aβ toxicity^63^. In cluster Crh/Gpc5 (27) of our dataset, gene sets related to Alzheimer’s Disease, Huntington’s Disease, Oxidative Phosphorylation, Parkinson’s Disease, Protein export, and Ribosome were all strongly under-enriched (Figure 7B), suggesting that this neuronal subtype has a distinct disease-associated expression signature compared to other neurons. Together, these data highlight for the first time the hypothalamic transcriptional changes unique to individual neuronal subtypes or common across neurons, which may contribute to age-related neurodegenerative disease.

Finally, we sought to understand the role of *Xist* in defining the aged neuronal state. To do so, we tested whether expression of X chromosome genes was sufficient to predict neuronal age in our dataset (Figure 8A). We trained eight different supervised machine learning models to classify neurons as either young or aged. Based on the accuracy score (Figure 8B), the XGBoost classifier (xgbc^64^) outperformed others with 77.8 ± 0.6224% accuracy. We then fine-tuned the model to optimize hyperparameters and retrained it on new data splits across 50 random states to measure the uncertainties due to splitting and the non-deterministic model. The confusion matrix (Figure 8C) and the area under the ROC curve (ROC AUC) (Figure 8D) confirmed model performance. Interestingly, when we randomly shuffled the feature *“Xist* expression”, the model performance dropped dramatically down to the near-baseline level (Figure 8E). We then applied Shapley additive explanations (SHAP)^65^ to further interpret the predictions and rank the features by importance. Consistent with our findings, *Xist* was the most important feature in the prediction, followed by the sum of all genes detected in the dataset (Figure 8F). Local feature importance of two randomly selected individual neurons (young and aged) also showed that *Xist* had the most significant impact on driving the model prediction (Figure 8G). These data suggest that *Xist* upregulation is a key feature of hypothalamic neuronal aging, and may predict female neuronal aging in the hypothalamus.

**Figure 8.**
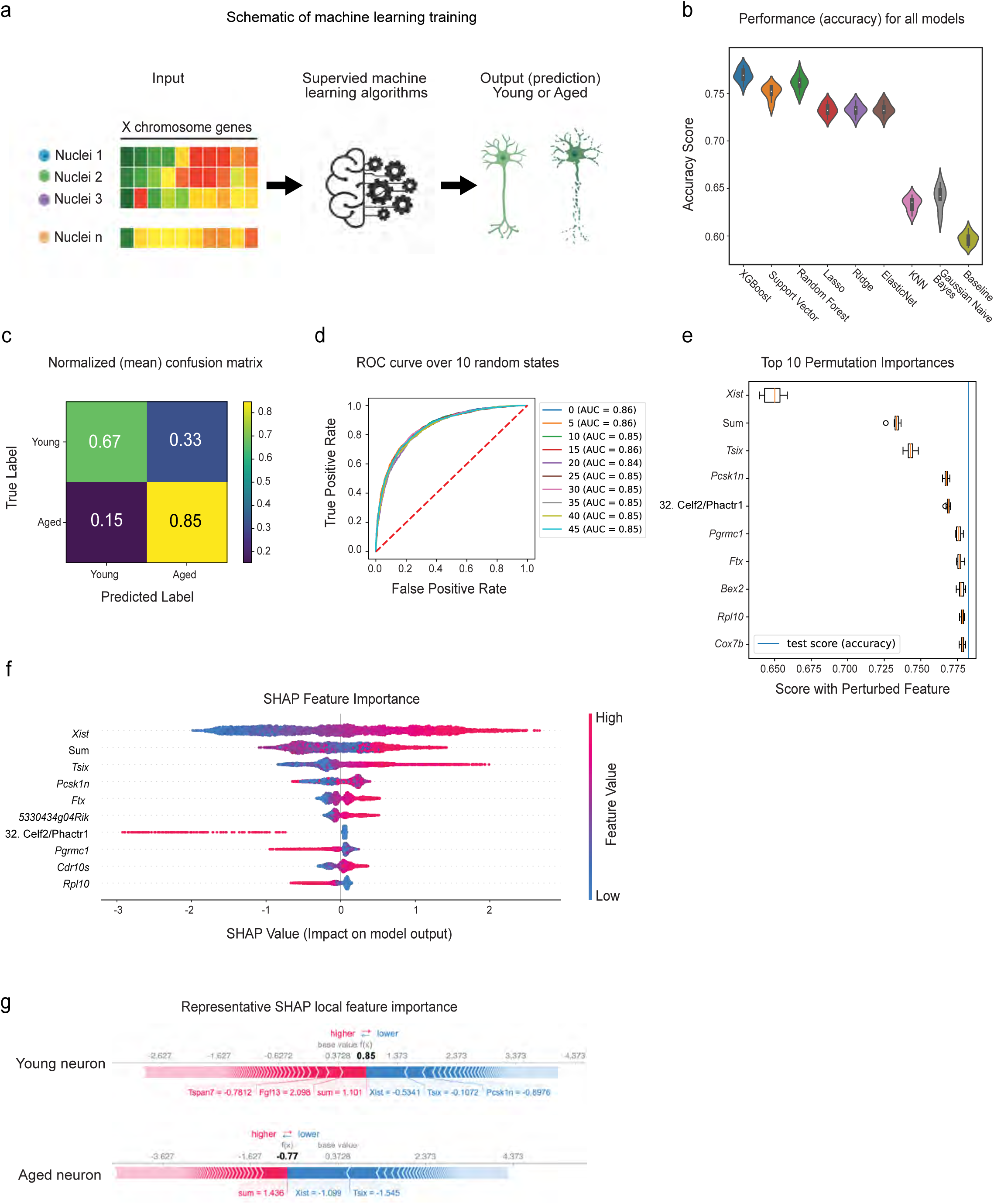
*Xist* expression predicts neuronal age in the mouse hypothalamus. A) Schematic of the machine learning approach. B) Violin plots showing model test set performance. Model accuracy across 10 random states is shown, with consistently XGBoost Classifier (xgbc) outperforming the other models. Xgbc was then retrained on new data splits across 50 different random states. C-D) Confusion matrix and ROC curve depicting Xgbc model accuracy across 50 and 10 random states respectively. E) Top 10 most important features of the Xgbc model. Note the strong influence of *Xist* on model accuracy score. F) SHAP summary plot showing feature importance for the top 10 features that predict cellular age in the model. G) SHAP force plot showing the most impactful features on the model prediction for example observations in young and aged neurons.

While comparing our dataset to hippocampal data, we noted that *Xist* is highly upregulated with age in both brain regions. To understand the role of *Xist* and the X chromosome in aging across brain regions, we tested whether X chromosome genes were sufficient to predict cellular age in the hippocampus. We reran the xbgc model using snRNA-seq data from female mouse hippocampal neurons (Figure 3A-Neuron, Supplementary Figure 6). Although overall gene expression changes with age in the hypothalamus and hippocampus do not correlate (Figure 2A), X chromosome gene expression is still sufficient to predict cellular age with 82.5 ± 0. 7080% accuracy in hippocampal neurons. Interestingly, *Xist* was the second most important predictor of age, based on permutation importance and SHAP (Supplementary Figure 6C-D), confirming *Xist* upregulation as a shared feature of neuronal aging across two brain regions.

## DISCUSSION

In this work, we used single nuclei RNA-seq to identify the age-associated transcriptional changes in the mouse hypothalamus. This brain region is critical for the regulation of physiological homeostasis, including sleep, circadian rhythms, feeding, and metabolism. These functions are well known to be disrupted during aging, and our findings implicate widespread transcriptional changes concomitant with physiological changes.

Our approach successfully captured the major cell types in the brain, as well as hypothalamus-specific cell-types such as tanycytes. We found that cellular subtypes in this region acquire distinct aging signatures, and discovered that increased transcriptional heterogeneity is a common feature across all cell types with age. Consistent with our findings, age-related transcriptional alterations have been observed in aging human brains and increased transcriptional noise is thought to be a hallmark of aging. Our finding that different neuronal subtypes have distinct aging signatures is consistent with recent reports identifying differential susceptibility to neurodegeneration^66^. Identification of the transcriptional signatures involved may pave the way for therapeutics targeted at subpopulations most susceptible to dysregulation with age.

We observed striking changes in the microglial population with age. Microglia are resident immune cells in the brain, and previous research has shown that microglia-mediated inflammation in the hypothalamus can affect lifespan^11^. By utilizing trajectory inference analysis, we uncovered that while some aging microglia retain features of young cells, the population shows a progression toward an aged phenotype based on distinct gene expression modules. Interestingly, disease-associated microglia genes such as *Apoe* change throughout both age and pseudotime.

Sex differences in aging have been observed across taxa, including in mice^12, 13^. In mammals, females generally live longer than males^12^, and many aging interventions such as CR, are more effective in females^13, 14^. In addition, the sexually dimorphic response to aging interventions appears to be a conserved phenomenon, with female *Drosophila* responding more strongly to dietary restriction paradigms than males^67^, and hermaphroditic *C. elegans* responding more strongly to DR than males^68^. In mice, males and females differ in regard to sex chromosome content (males are XY and females are XX) and the presence of gonadal hormones such as higher androgens in males and estrogens in females. Interestingly, X chromosome content has been linked to longevity, and the presence of two X chromosome contributes to increased longevity regardless of hormonal status^69^. This study from the Dubal lab was performed using the four core genomes mouse line, in which the *Sry* gene (which induced testes development) exists on an autosome rather than the Y chromosome, allowing for chromosomal sex to be disambiguated from gonadal sex/hormone status. In our study, we uncover a potential mechanism by which the X chromosome affects aging. We observed widespread upregulation of *Xist* in aged female animals, as well as upregulation of other XIC genes including *Tsix*, *Jpx*, and *Ftx*. Intriguingly, this increased expression was highly prominent in neurons, although upregulation of *Xist* in was observed in oligodendrocytes, astrocytes, and tanycytes as well. Strikingly, in a machine learning algorithm, *Xist* expression was the most important variable in a model to classify whether a hypothalamic neuron was young or aged. Together, our findings reveal a novel feature of aging in the female brain. Moreover, recent research suggests a general role for long non-coding RNAs in the maintenance of chromatin structure^70^, and previous single cell RNA sequencing studies of the aged brain identify upregulation of long non-coding RNAs such as *Malat1* as a feature of aging^37^. Together, this work suggests that that understanding the mechanisms and consequences of *Xist* upregulation in aging may provide novel insight into sex differences in aging.

In summary, our study reveals the major transcriptional features of hypothalamic aging. We observed transcriptional variation across cell types, cell-type specific aging signatures, and novel features of aging in females. Understanding how individual populations of cells in this region contribute to overall loss of homeostasis with age will be vital to identifying treatments for aging and age-related disease.

## METHODS

### Animals

#### Single nuclei isolation

Young (3 month) and aged (19-24 month) C57/Bl6 female mice were obtained from the National Institute on Aging. Mice were housed and used according to protocols approved by Brown University and in accordance with institutional and national guidelines. Animals were exposed to male bedding 3 days before sacrifice to sync estrous cycle. Animals were sacrificed at Zeitgeiber time ZT2-ZT3.

### Single nuclei RNA-sequencing

In order to reduce noise stemming from differences in estrous state, two whole hypothalamuses were pooled into each biological replicate, for a total of two replicates for the young and aged conditions. Nuclei extraction was performing using the Nuclei Isolation Kit: Nuclei PURE Prep Kit (Millipore Sigma) according to the manufacturer’s instructions with the following modifications: for each sample, two hypothalamuses were dissected out of the animals and rinsed in cold PBS. Tissue was transferred using a transfer pipette into a refrigerated Dounce homogenizer with 5 mL of lysis solution following kit instructions. Tissue was homogenized with the Dounce B and the lysate was transferred into a 15 mL falcon tube through a 40-micron filter. The sucrose purification step was performed with the following modifications: half the volume of all reagents was used to account for the small tissue sample sizes, an SW34 rotor was used, and samples were spun for 45 minutes at 30,000 X g (13,000 rpm) at 4 °C. Nuclei were counted using a hemocytometer, and 5000 cells per sample were loaded onto the Chromium Single Cell 3′ Chip (10x Genomics) and processed with the Chromium Controller (10x Genomics). Samples Young _1, Young_2, Aged_1 and Aged_2 were prepared using the Chromium Single Cell 3′ Library & Gel Bead kit v2 according to manufacturer’s instructions. Samples were sequenced at GENEWIZ, Inc on an Illumina HiSeq, with a target of 50,000 reads per sample. The Aged_1 and Young_2 samples underwent an additional round of sequencing to obtain sufficient read depth. Samples Young_3, Young_4, Aged_3, and Aged_4 were prepared with the Next GEM Single Cell 3ʹ Reagent kit (10x Genomics) and sequenced at Genewiz on an Illumina NovaSeq.

### Quality control, data processing and analysis

We performed sequence alignment to the mm10 genome (2020) using the CellRanger (cellranger/6.0.0) software from 10x Genomics with the –include introns flag. The resulting feature-barcode matrices were read into R version 4.1.0, excluding any nuclei expressing fewer than 200 genes, and any gene expressed in fewer than three nuclei.

Filtering and visualization were performed using Seurat (4.0.3)^71^. For samples sequenced on an Illumina HiSeq, nuclei with fewer than 200 or more than 3000 features were filtered out. For samples sequenced on the NovaSeq, nuclei with fewer than 200 or more than 7500 features were filtered out. Similarly, nuclei with greater than 10% mitochondrial mapping were removed, resulting in 23,808 nuclei in the aged condition, and 16,256 nuclei in the young condition. Integration of the datasets was performed using the IntegrateData function on 5000 variable features. The number of nuclei, unique molecular identifiers, and unique genes per sample are reported in Supplementary Figure 1. To assign identities to clusters, the FindAllMarkers() command with default parameters was used. This finds the top genes that define a cluster identity. We named each cluster using the top 2 genes to come out of the FindAllMarkers() analysis.

Differential expression was performed using MAST(1.18.0)^30, 31^, with random effect for sequencing depth and sample of origin^69^. Genes were considered significant if the adjusted p-value was less than 0.05, and the log2 fold change was greater than 0.1 or less than -0.1. For re-analysis of publicly available data, raw cell/count matrices were downloaded, and data was reprocessed according to above workflow. MAST was performed with random effect for sample of origin.

### Gene Set Enrichment Analysis

Gene Set Enrichment Analysis was performed using the fgsea package (1.18.0)^38^ using the Hallmark gene set list and KEGG gene set list from MSigDB (version 7.2.)^72^. For each cluster, genes were ranked by log2 fold change after MAST analysis, and the analysis was performed using the fgseaMultilevel command with default settings and seed set at 1000. Gene sets were considered to be enriched if the adjusted p value was less than 0.1. Conversions between mouse and human annotation was performed using biomaRt (2.48.2).

### Trajectory inference and analysis using Monocle3

To infer the aging process for the microglia/macrophage clusters (n = 1121 nuclei) generated in Seurat, we applied Monocle3^46, 59^. Monocle3 uses dimensionality reduction to place single cells in a 2D space, removes batch effects by mutual nearest neighbor alignment, and connects single cells to construct a trajectory in a semi-supervised way. For the microglia/macrophage cluster, we use the integrated Seurat object with no further batch correction or dimensionality reduction in Monocle3. We subsetted the microglia and macrophage cluster and programmatically specified the root of the trajectory by selecting the node most enriched for young cells. The trajectory and its direction calculated by Monocle3 are in agreement with the distribution of young and aged cells. Spatially differential expression analysis along the trajectory was performed with Moran’s I test in Monocle3, and selected genes with q < 0.05 as trajectory-dependent genes (2112 genes). The set of genes were grouped into four modules according to its RVAE decoded expression^49^ along the trajectory.

### Functional enrichment analysis

enrichR^73^ 3.0 was applied to perform the functional enrichment analysis of 2112 genes in individual modules, resulting in lists (“4_modules_q_moranI”) of statistically significant enriched terms (adjusted p < 0.05 with Benjamini-Hochberg correction) for individual modules. We checked the gene sets database GO_Biological_Process_2018, GO_Cellular_Component_2018, and GO_Molecular_Function_2018. We kept GO terms with p < 0.05, and visualized the 10 most significant terms for each module and visualized in the dotplot. The python package RVAgene (version 1.0, in python version 3.9.6 with PyTorch version 1.9.0) recurrent variational autoencoder (RVAE) implementation was used to decode the trajectory-differentially expressed genes (t-DEGs, n=2112) along the pseudotime trajectory. Expression was averaged in individual time bins, and then rescaled to the value in [-1,1] and input to RVAgene. For the neural network, the following parameters were used: symmetrical architecture with two hidden layers (48 nodes per layer) and two latent variable dimensions. The output reconstructed trajectory for the t-DEGs was used to plot the heatmap.

### Single-Cell rEgulatory Network Inference and Clustering (SCENIC)

RNA counts from samples Young_3, Young_4, Aged_3 and Aged_4 were exported into a loom file using SCopeLoomR_0.11.0. The standard pySCENIC workflow was run using Brown University’s cloud computing resource. The workflow was completed 50 times, and the resulting loom files were loaded back into R. Only regulons and genes within the regulons appearing 10 out of 50 times or more were retained for further analysis. AUCell analysis was performed in R using SCENIC_1.2.4 and AUCell_1.14.0. Using the previously defined regulons, AUCell analysis was performed on all cells from the dataset following the default pipeline. For binarization of regulons, default thresholds were used. Regulon specificity scores were generated using the calcRSS() command.

### CellPhoneDB

RNA counts for young and aged datasets were analyzed separately to allow for comparison. Conversions between mouse and human annotation was performed using biomaRt (2.48.2). CellPhoneDB (version 2.1.7) was run in a conda environment (anaconda/2020.02) using the statistical_analysis method with 1000 iterations and a .1 threshold. For visualization in R, only ligand-receptor pairs in which direction could be inferred were retained for analysis.

### Microglia isolation and RT-qPCR analysis

Young (2-3 month), middle aged (8-13 month), and aged (20-24 month) C57/Bl6 wild type and POMC-EGFP reporter mice (Jax Stock No: 009593) were housed and used according to protocols approved by Brown University and in accordance with institutional and national guidelines. Animals were sacrificed at ZT4. For each biological replicate, four animals were pooled, with genotypes and estrous state balanced across conditions. Tissue was dissociated with the Adult Brain Dissociation Kit (Miltenyi Biotec, # 130-107-677) according to manufacturer’s instructions. Dissociated tissue was incubated CD11b MicroBeads (Miltenyi Biotec, #130-049-601) for fifteen minutes at 4^0^C. Labeled cells were isolated using Miltenyi Biotec MS columns (#130-042-201) on the OctoMACS Separator. RNA was purified using the RNeasy micro kit (#74004) and cDNA was generated with the High-Capacity Reverse Transcription Kit (Applied Biosystems #4374966). A negative control (-RT) for each sample was also generated by excluding the Multiscribe Reverse Transcriptase component of the reaction.

### Cell2location

Cell2location is a Bayesian model that uses snRNAseq cell type signatures to infer cell types in Visium spatial transcriptomics by decomposing mRNA counts in each Visium voxel into cell types. We performed the three main steps in the cell2location workflow: estimate reference expression signatures of cell types using our dataset, map the learned cell type signatures onto the slides, and performed downstream analysis including Pearson correlation, visualization, and NMF. The code, model parameters and training evaluations can be found in the jupyter notebooks in our github repository. In brief, we used the default parameters to train the cell2location model.

### Neuronal age prediction using machine learning

Neuronal nuclei (25,002) were selected for young or aged classification. All genes were annotated with their chromosomal location. For each neuron, one categorical feature (neuronal subtype) and 281 numerical features were used for machine learning: 278 X chromosome genes (mean expression > 0.1 read per cell and detected in > 3,000 nuclei), aggregated X chromosome gene expression “x_sum”, aggregated all gene expression “sum”, and their ratio “x_prop”. The pipeline and functions were implemented in Scikit-learn^74^. For data splitting, 20% of nuclei were first split into the testing set, and the rest 80% were further split into training and validation sets using 5 fold cross-validation, resulting in train-validation-testing = 64-16-20. For preprocessing, OneHotEncoder was applied for the categorical feature, and StandardScaler was applied for the numerical features.

**Table 1.** Parameters used for tuning models

| Model | Parameters |
| --- | --- |
| L1 (Lasso) | C: 0.01, 0.05, **0.1**, 0.5, 1, 5, 10 |
| L2 (Ridge) | C: 0.01, 0.05, **0.1**, 0.5, 1, 5, 10 |
| ElasticNet | C: 0.05, **0.1**, 0.5, 1, 3; **l1_ratio**: 0.2, 0.35, **0.5**, 0.65, 0.8 |
| Random Forest | **max_features**: 25, 50, **75**, 100, 200, None; **max_depth**: 10, 20, **30**, 50, 100, None; **min_samples_split**: 2, 5, **10**, 20 |
| SVC | **gamma**: 1e-4, 1e-3, **1e-2**, 1e-1; C: np.logspace(-1, 1, 5) (**3.162**) |
| XGBoost | **max_depth**: 1, 3, **5**, 10, 20, 30, 100; 1, 2, 3, 4, **5**, 8, 10, 15, 20 (finer) |
| KNN | **n_neighbors**: 30, 100, 200, 300; **weights**: uniform, distance |

SVC: Support Vector Machine Classifier with rbf kernel,
KNN: K Nearest Neighbor
The best parameters are bolded or indicated next to the grid.

Eight models (above) were tested over ten different random states. The best hyperparameters were selected using GridSearchCV, and the model performance was evaluated using accuracy score of the test sets. XGBoost Classifier^64^ was selected, fine-tuned (max_depth = 5 with early stop), and then retrained on new splits across 50 different random states. The baseline accuracy was 0.596 ± 0.00765, while the model accuracy was 0.778 ± 0.006224. Model interpretation was performed using permutation feature importance and SHAP^65^.

For the hippocampus dataset, neuronal nuclei (11,204) were selected. For each neuron, 253 X chromosome genes (mean expression > 0.1 read per cell and detected in > 1,000 nuclei), aggregated X chromosome gene expression “x_sum”, aggregated all gene expression “sum”, and their ratio “x_prop” were used as features for model training. The rest of the processes were the same as above except that the max_depth = 4 for the final 50 different random states. The baseline accuracy was: 0.594 ± 0.009167, while the model accuracy was : 0.825 ± 0.007080.

### Data and code availability

Fastq files for raw single nuclei RNA sequencing and Seurat object were deposited at GEO accession XYZ. Code available https://github.com/Webb-Laboratory/Hajdarovic_And_Yu_et_al_2022. Publicly available datasets are available on GEO: Hippocampal single nuclei data, GSE161340; human entorhinal cortex data, GSE138852 (samples AD3-AD4 and Ct1-Ct2); human prefrontal cortex data, GSE174367 (samples 17, 19, 37, 43, 45, 50, 66, 90); Spatial data, ArrayExpress E-MTAB-11114.

### Whole brain RNA isolation and cDNA generation and qRT-PCR

Hypothalamus, olfactory bulb, cerebellum, and cortex were dissected in cold PBS from the brains of 3-month-old and 24 month old C57Bl/6 mice (n=6, 3 male and 3 female for each age) and snap frozen in liquid nitrogen. RNA was purified using the Qiagen RNeasy Lipid Tissue Mini Kit (Qiagen #74804). cDNA was generated using 500 ng of RNA and the High-Capacity Reverse Transcription Kit (Applied Biosystems #4374966). A negative control (-RT) for each sample was also generated by excluding the Multiscribe Reverse Transcriptase component of the reaction. qPCR reactions were completed using the PowerUp^TM^ SYBR ^TM^ Green Master Mix (Invitrogen #A25918). Stock primers were diluted to 10 mM in sterile water, and cDNA was diluted 1:5 in sterile water (whole brain) or 1:3 in sterile water (microglia). Expression levels of the genes of interest (see table below) were quantified using a ViiA 7 Real Time PCR System with QuantStudio software. For whole brain, *Actin* was used as a housekeeping gene. For microglia, *Itgam* (CD11b) was used. Each sample, water control, and - RT control sample was run in triplicate for each primer set. CT values were normalized to the housekeeping gene, and ΔCT values were plotted as 2^-ΔCT^. Technical replicates were averaged per biological replicate.

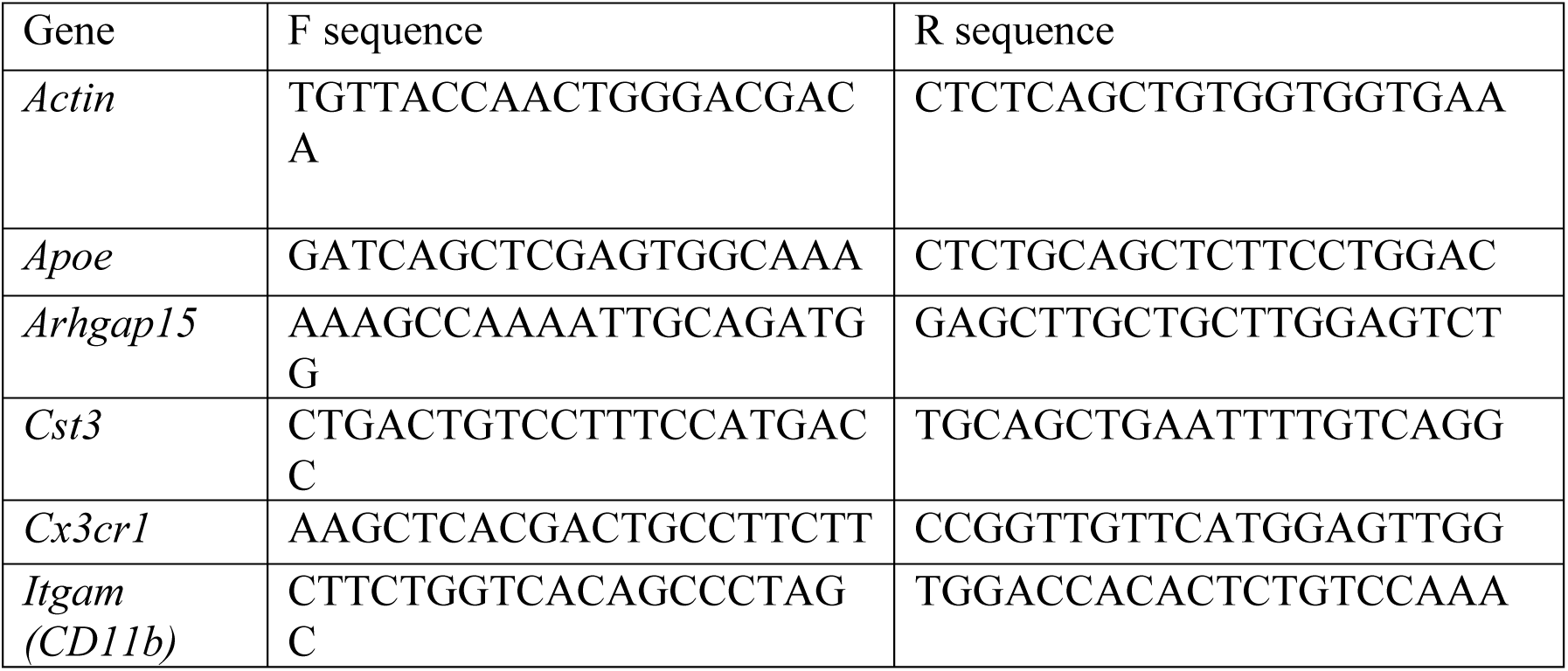
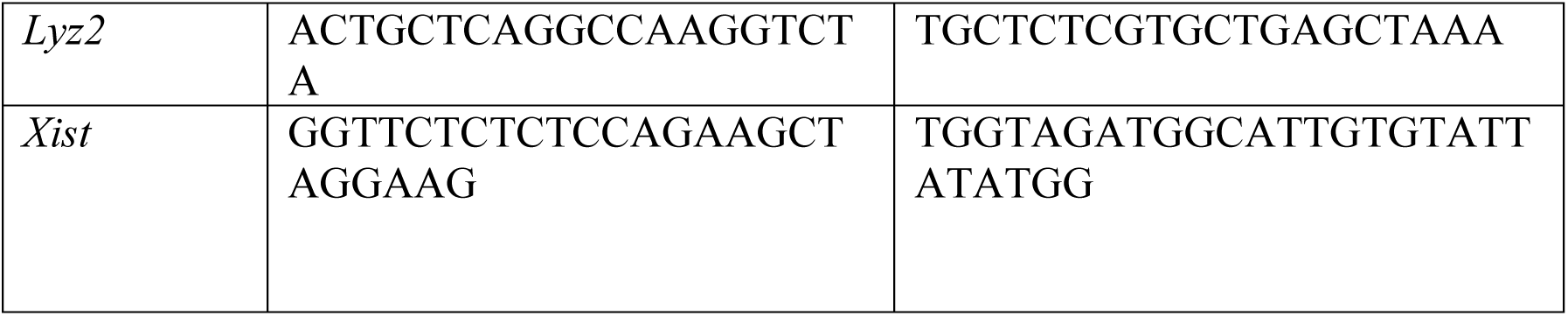

## SUPPLEMENTARY MATERIAL

Supplementary Table 1. Markers for hypothalamic cell clusters.

Supplementary Table 2. Binarized regulons for hypothalamic cell clusters.

Supplementary Table 3. Results of differential expression analysis for all nuclei.

Supplementary Table 4. Differential expression analysis of individual cell types.

Supplementary Table 5. Results of Moran’s I Test.

Supplementary Table 6. Cluster markers for neuronal subtypes.

Supplementary Table 7. Neuropeptide genes in each cluster and function of gene

Supplementary Table 8. Results of differential expression for neuronal subtypes.

**Supplementary Table 7.**
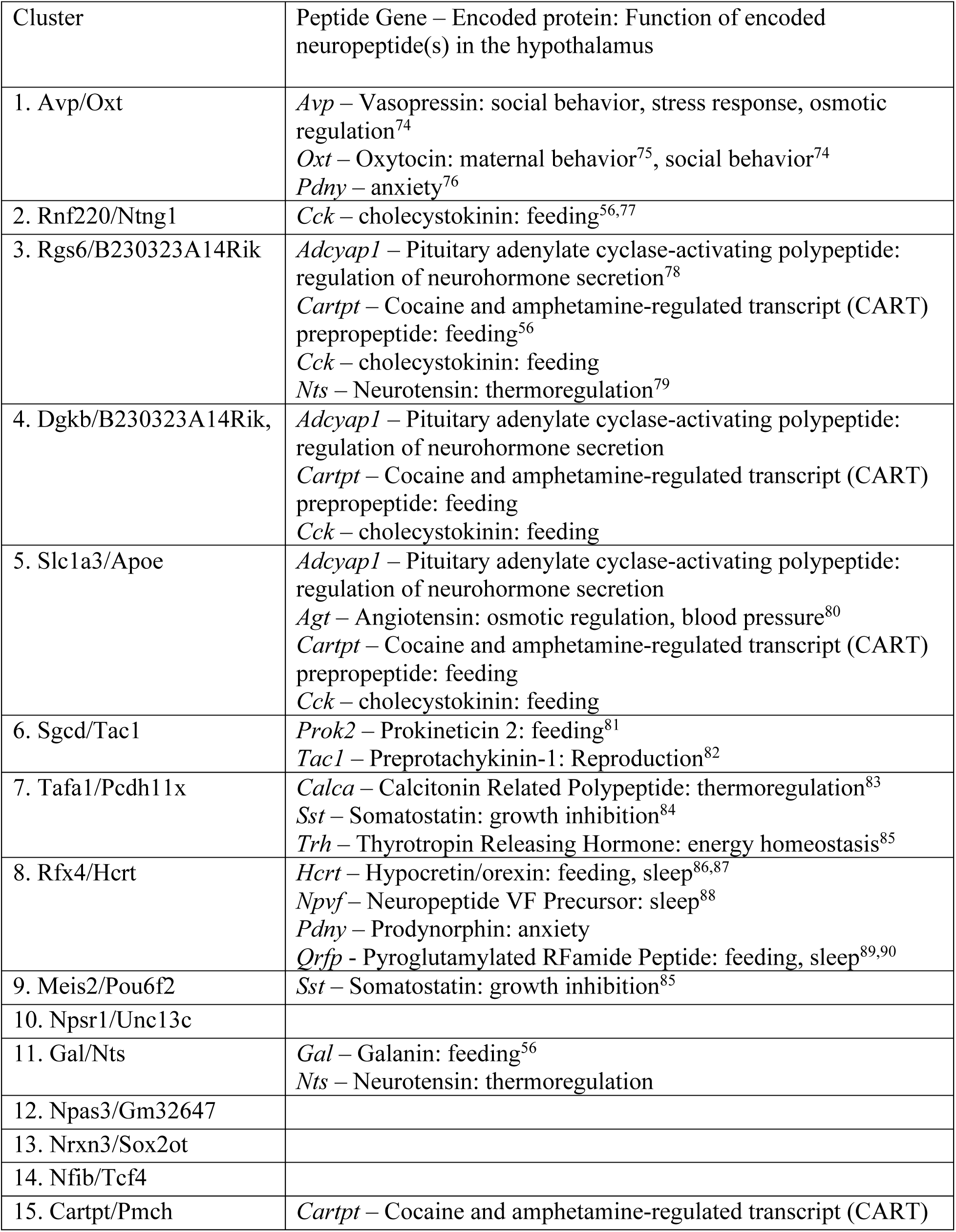
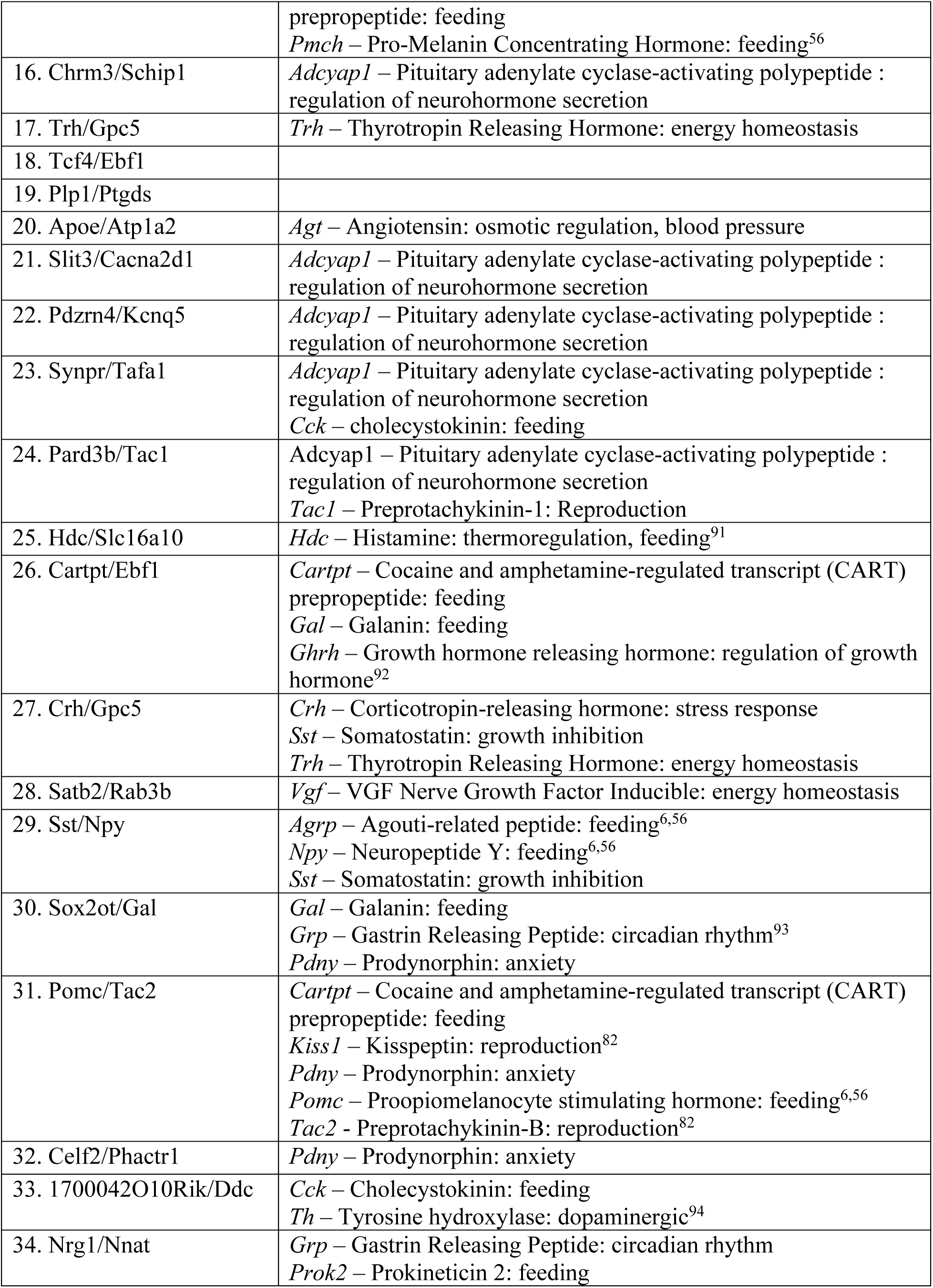
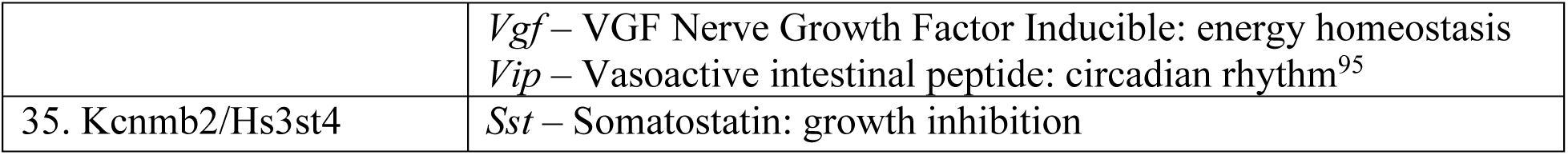

---

**Supplementary Figure 1.**
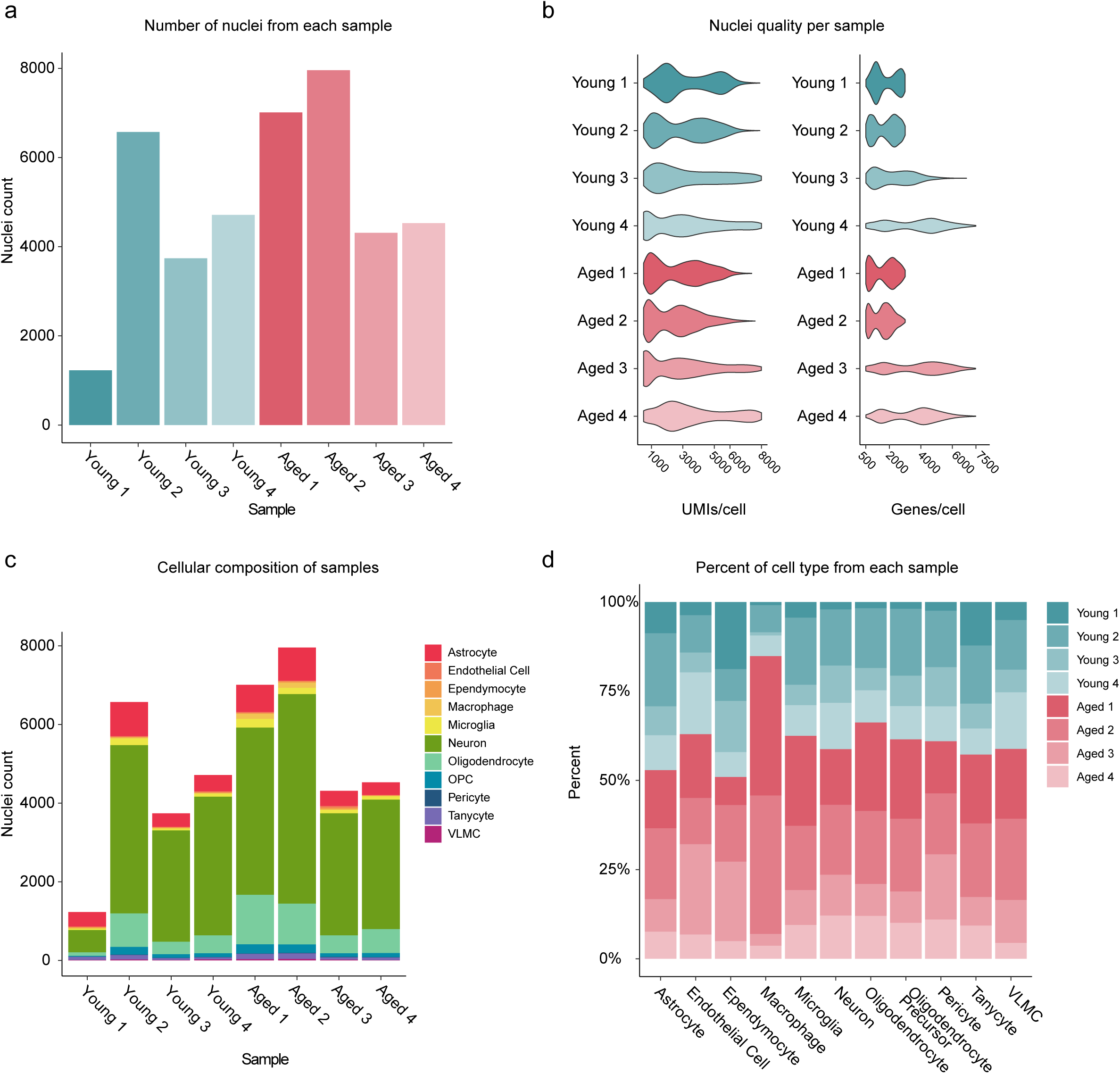
Quality control metrics for single nuclei data. A) Number of nuclei per sample. B) Violin plots showing the number of UMIs per nuclei per sample (left) and the number of genes per nuclei per sample (right). C) Number of nuclei assigned to a cellular subtype per sample. D) Proportion of each cell type arising from a particular sample.

**Supplementary Figure 2.**
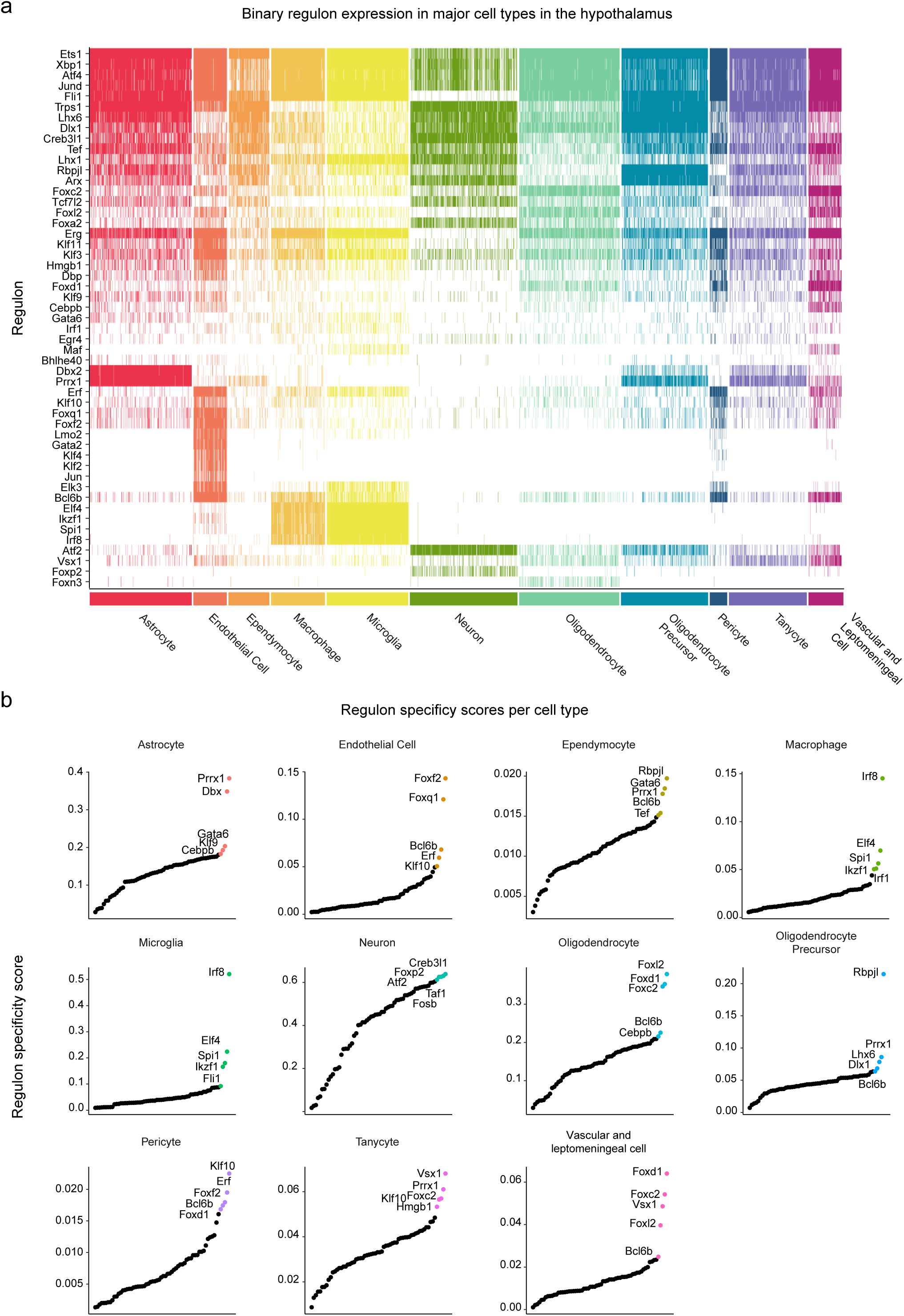
Gene regulatory network reconstruction for cellular subtypes. A) Binarized regulon activity for each regulon in each nuclei. Maximum 500 nuclei per cluster shown. Color indicates a regulon is “on” in each cell. B) Regulon specificity score indicating whether a given regulon is specific to a cell type. Top 5 RSS for each cluster shown.

**Supplementary Figure 3.**
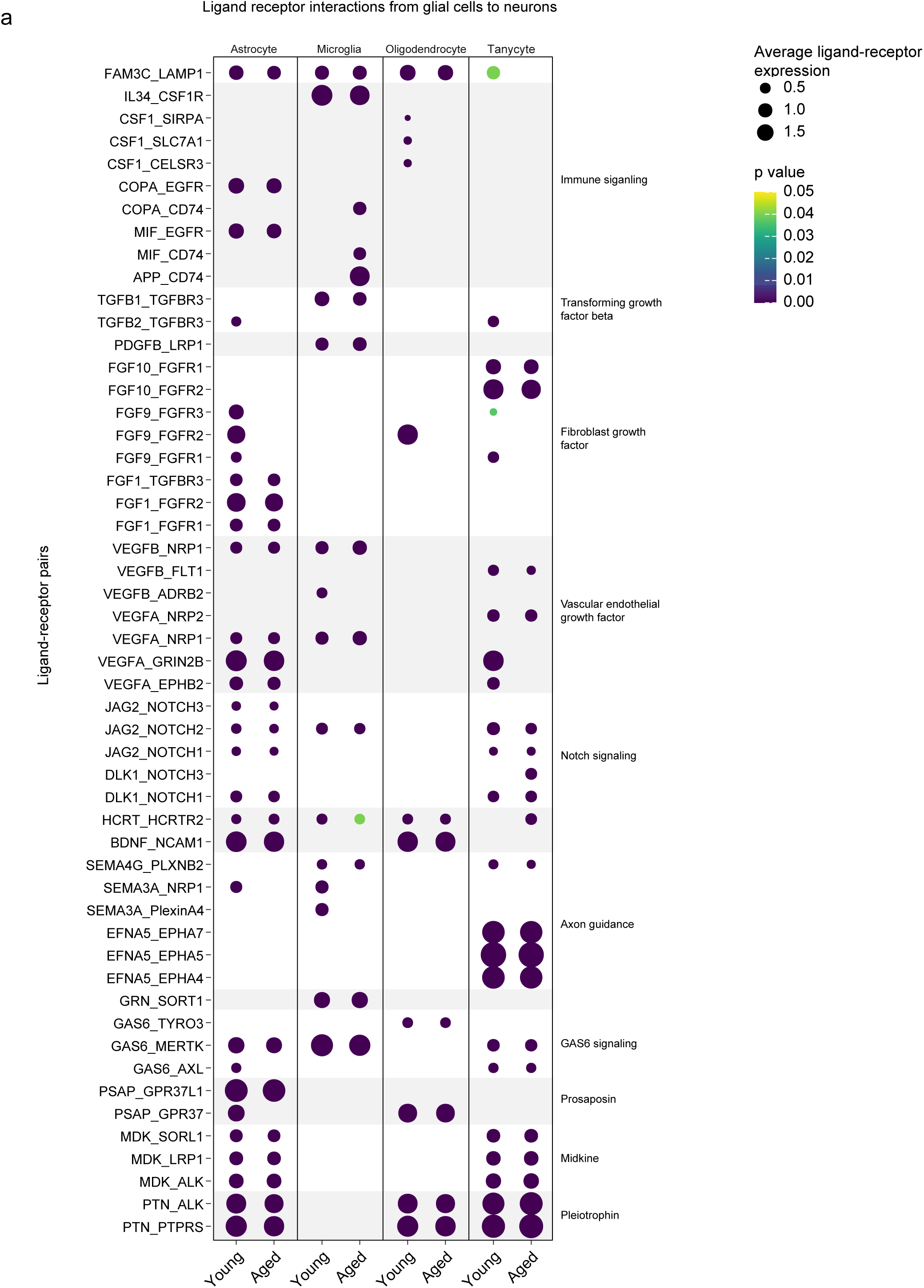
Non-neuronal signaling to neurons in young and aged hypothalamus. For each ligand-receptor pair, dot size equals mean expression of ligand gene in signaling cell and receptor gene in receiving cell, color equals p value.

**Supplementary Figure 4.**
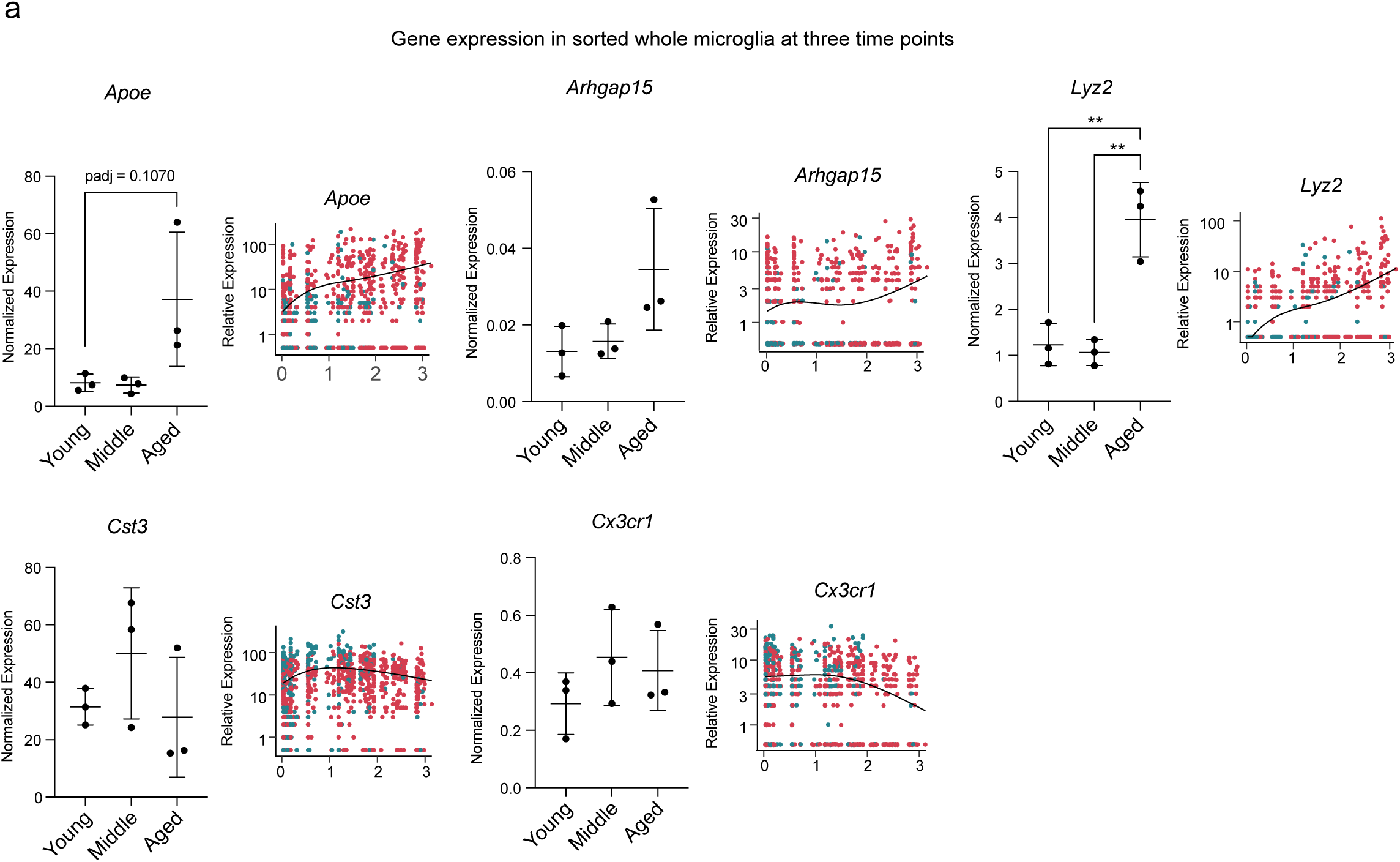
RT-qPCR of sorted whole CDllb+ microglia. One way ANOVA with Holm-Šídák test, **, adjusted p value < 0.01). Shown with pseudotime trajectories from figure 4.

**Supplementary Figure 5.**
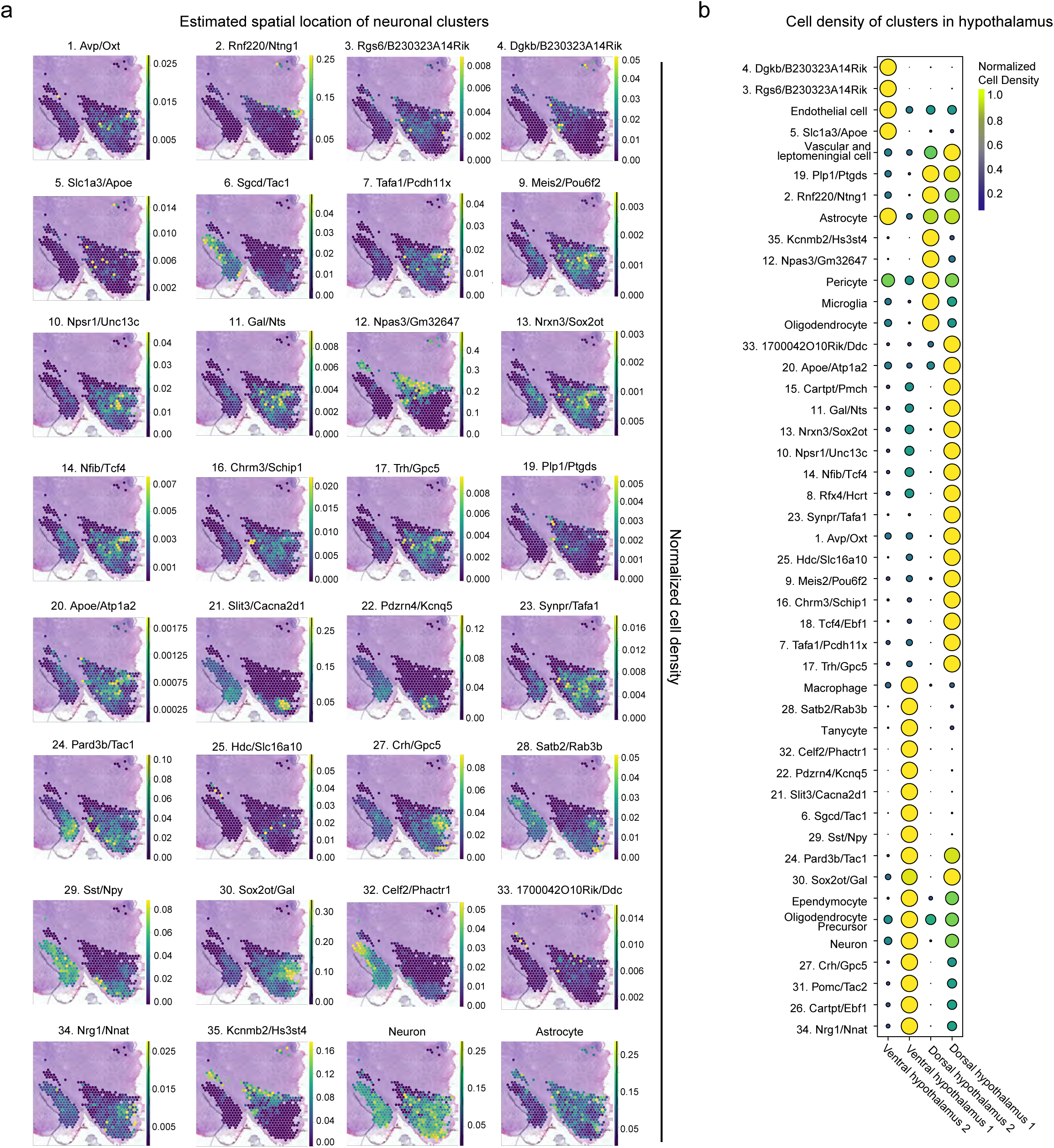
A) Normalized estimated cell density of neuronal subclusters in coronal section of spatial transcriptomic data. B) Estimated cell density of broad cell categories and neuronal subclusters in total spatial transcriptional data.

**Supplementary Figure 6.**
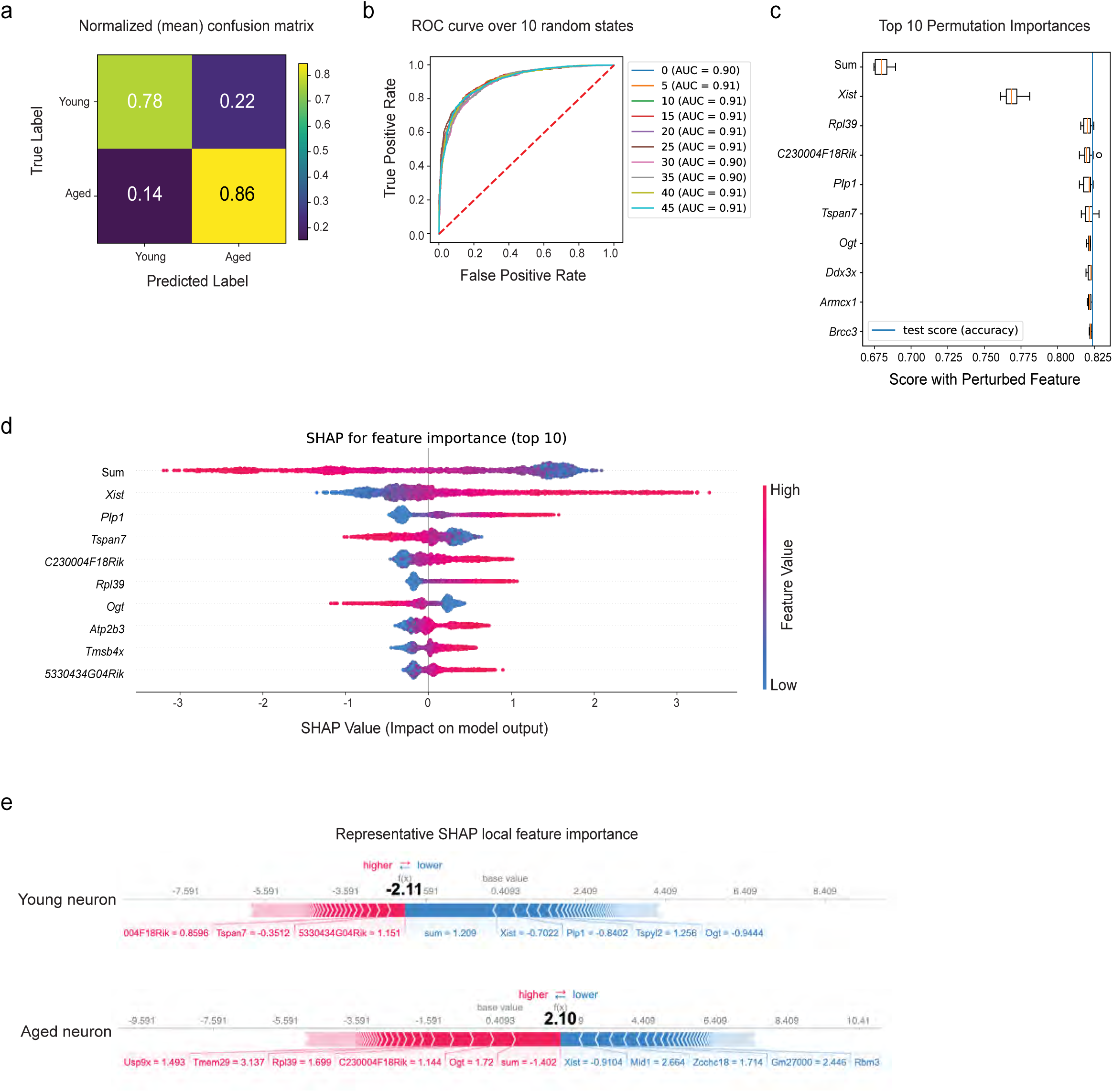
*Xist* expression predicts neuronal age in the mouse hippocampus. A-B) Confusion matrix and ROC curve depicting Xgbc model accuracy across 50 and 10 random states, respectively. C) Top 10 most important features of the Xgbc model. Note the strong influence of *Xist* on model accuracy score. D) SHAP summary plot showing feature importance for the top 10 features that predict cellular age in the model. E) SHAP force plot showing the most impactful features on the model prediction for example observations in young and aged neurons.

## BIBLIOGRAPHY.

1. Partridge, L., Deelen, J. & Slagboom, P. E. Facing up to the global challenges of ageing. Nature 561, 45–56 (2018).

2. Yousufuddin, M. & Young, N. Aging and ischemic stroke. Aging 11, 2542–2544 (2019).

3. Balducci, L. & Aapro, M. Epidemiology of Cancer and Aging. in Biological Basis of Geriatric Oncology 1–15 (Springer US, 2005).

4. Farooqui, T. & Farooqui, A. A. Aging: An important factor for the pathogenesis of neurodegenerative diseases. Mech. Ageing Dev. 130, 203–215 (2009).

5. Chahal, H. & Drake, W. The endocrine system and ageing. J. Pathol. 211, 173–180 (2007).

6. Toda, C., Santoro, A., Kim, J. D. & Diano, S. POMC Neurons: From Birth to Death. Annu. Rev. Physiol. 79, 209–236 (2017).

7. de Cabo, R., Carmona-Gutierrez, D., Bernier, M., Hall, M. N. & Madeo, F. The Search for Antiaging Interventions: From Elixirs to Fasting Regimens. Cell 157, 1515–1526 (2014).

8. Bishop, N. A. & Guarente, L. Two neurons mediate diet-restriction-induced longevity in C. elegans. Nature 447, 545–549 (2007).

9. Broughton, S. J. et al. DILP-producing Median Neurosecretory Cells in the Drosophila Brain Mediate the Response of Lifespan to Dietary Restriction. Aging Cell 9, 336–346 (2010).

10. Satoh, A. et al. Sirt1 extends life span and delays aging in mice through the regulation of Nk2 homeobox 1 in the DMH and LH. Cell Metab. 18, 416–430 (2013).

11. Zhang, G. et al. Hypothalamic programming of systemic ageing involving IKK-β, NF-κB and GnRH. Nature 497, 211–216 (2013).

12. Lemaître, J.-F. et al. Sex differences in adult lifespan and aging rates of mortality across wild mammals. Proc. Natl. Acad. Sci. U. S. A. 117, 8546–8553 (2020).

13. Austad, S. N. & Bartke, A. Sex Differences in Longevity and in Responses to Anti-Aging Interventions: A Mini-Review. Gerontology 62, 40–46 (2016).

14. Mitchell, S. J., et al. Effects of Sex, Strain, and Energy Intake on Hallmarks of Aging in Mice. Cell Metab. 23, 1093–1112 (2016).

15. Satoh, A. et al. SIRT1 Promotes the Central Adaptive Response to Diet Restriction through Activation of the Dorsomedial and Lateral Nuclei of the Hypothalamus. J. Neurosci. 30, 10220– 10232 (2010).

16. Webb, A. E., Kundaje, A. & Brunet, A. Characterization of the direct targets of FOXO transcription factors throughout evolution. Aging Cell 15, 673–685 (2016).

17. Benayoun, B. A. et al. Remodeling of epigenome and transcriptome landscapes with aging in mice reveals widespread induction of inflammatory responses. Genome Res. 29, 697–709 (2019).

18. Hofmann, J. W. et al. Reduced Expression of MYC Increases Longevity and Enhances Healthspan. Cell 160, 477–488 (2015).

19. Chen, R., Wu, X., Jiang, L. & Zhang, Y. Single-Cell RNA-Seq Reveals Hypothalamic Cell Diversity. Cell Rep. 18, 3227–3241 (2017).

20. Mickelsen, L. E. et al. Single-cell transcriptomic analysis of the lateral hypothalamic area reveals molecularly distinct populations of inhibitory and excitatory neurons. Nat. Neurosci. 22, 642–656 (2019).

21. Campbell, J. N. et al. A Molecular Census of Arcuate Hypothalamus and Median Eminence Cell Types. Nat. Neurosci. 20, 484–496 (2017).

22. Mickelsen, L. E. et al. Neurochemical Heterogeneity Among Lateral Hypothalamic Hypocretin/Orexin and Melanin-Concentrating Hormone Neurons Identified Through Single-Cell Gene Expression Analysis. eNeuro 4, (2017).

23. Mickelsen, L. E. et al. Cellular taxonomy and spatial organization of the murine ventral posterior hypothalamus. eLife 9, e58901 (2020).

24. Kim, D. W. et al. The cellular and molecular landscape of hypothalamic patterning and differentiation from embryonic to late postnatal development. Nat. Commun. 11, 4360 (2020).

25. Romanov, R. A. et al. Molecular design of hypothalamus development. Nature 582, 246–252 (2020).

26. Wen, S. et al. Spatiotemporal single-cell analysis of gene expression in the mouse suprachiasmatic nucleus. Nat. Neurosci. 23, 456–467 (2020).

27. Bakken, T. E. et al. Single-nucleus and single-cell transcriptomes compared in matched cortical cell types. PLOS ONE 13, (2018).

28. Ding, J. et al. Systematic comparison of single-cell and single-nucleus RNA-sequencing methods. Nat. Biotechnol. 38, 737–746 (2020).

29. Van de Sande, B. et al. A scalable SCENIC workflow for single-cell gene regulatory network analysis. Nat. Protoc. 15, 2247–2276 (2020).

30. Finak, G. et al. MAST: a flexible statistical framework for assessing transcriptional changes and characterizing heterogeneity in single-cell RNA sequencing data. Genome Biol. 16, 278 (2015).

31. Zimmerman, K. D., Espeland, M. A. & Langefeld, C. D. A practical solution to pseudoreplication bias in single-cell studies. Nat. Commun. 12, 738 (2021).

32. Jiang, C. H., Tsien, J. Z., Schultz, P. G. & Hu, Y. The effects of aging on gene expression in the hypothalamus and cortex of mice. Proc. Natl. Acad. Sci. U. S. A. 98, 1930–1934 (2001).

33. Arsenijevic, Y., Dreifuss, J. J., Vallet, P., Marguerat, A. & Tribollet, E. Reduced binding of oxytocin in the rat brain during aging. Brain Res. 698, 275–279 (1995).

34. Brockdorff, N. & Duthie, S. M. X chromosome inactivation and the Xist gene. Cell. Mol. Life Sci. CMLS 54, 104–112 (1998).

35. Lee, J. T., Davidow, L. S. & Warshawsky, D. Tsix, a gene antisense to Xist at the X-inactivation centre. Nat. Genet. 21, 400–404 (1999).

36. Ogrodnik, M. et al. Whole-body senescent cell clearance alleviates age-related brain inflammation and cognitive impairment in mice. Aging Cell 20, e13296 (2021).

37. Ximerakis, M. et al. Single-cell transcriptomic profiling of the aging mouse brain. Nat. Neurosci. 22, 1696–1708 (2019).

38. Sergushichev, A. A. An algorithm for fast preranked gene set enrichment analysis using cumulative statistic calculation. bioRxiv 060012 (2016) doi:10.1101/060012.

39. Boisvert, M. M., Erikson, G. A., Shokhirev, M. N. & Allen, N. J. The Aging Astrocyte Transcriptome from Multiple Regions of the Mouse Brain. Cell Rep. 22, 269–285 (2018).

40. Koseoglu, M. M., Norambuena, A., Sharlow, E. R., Lazo, J. S. & Bloom, G. S. Aberrant Neuronal Cell Cycle Re-Entry: The Pathological Confluence of Alzheimer’s Disease and Brain Insulin Resistance, and Its Relation to Cancer. J. Alzheimers Dis. JAD 67, 1–11 (2019).

41. González-García, I. et al. mTOR signaling in the arcuate nucleus of the hypothalamus mediates the anorectic action of estradiol. J. Endocrinol. 238, 177–186 (2018).

42. Efremova, M., Vento-Tormo, M., Teichmann, S. A. & Vento-Tormo, R. CellPhoneDB: inferring cell–cell communication from combined expression of multi-subunit ligand–receptor complexes. Nat. Protoc. 15, 1484–1506 (2020).

43. Reuss, B. & von Bohlen und Halbach, O. Fibroblast growth factors and their receptors in the central nervous system. Cell Tissue Res. 313, 139–157 (2003).

44. Baldwin, K. T. & Eroglu, C. Molecular mechanisms of astrocyte-induced synaptogenesis. Curr. Opin. Neurobiol. 45, 113–120 (2017).

45. Luo, X.-G., Ding, J.-Q. & Chen, S.-D. Microglia in the aging brain: relevance to neurodegeneration. Mol. Neurodegener. 5, 12 (2010).

46. Trapnell, C. et al. The dynamics and regulators of cell fate decisions are revealed by pseudotemporal ordering of single cells. Nat. Biotechnol. 32, 381–386 (2014).

47. Deczkowska, A. et al. Disease-Associated Microglia: A Universal Immune Sensor of Neurodegeneration. Cell 173, 1073–1081 (2018).

48. Keren-Shaul, H. et al. A Unique Microglia Type Associated with Restricting Development of Alzheimer’s Disease. Cell 169, 1276–1290.e17 (2017).

49. Mitra, R. & MacLean, A. L. RVAgene: generative modeling of gene expression time series data. Bioinformatics 37, 3252–3262 (2021).

50. Chureau, C. et al. Ftx is a non-coding RNA which affects Xist expression and chromatin structure within the X-inactivation center region. Hum. Mol. Genet. 20, 705–718 (2011).

51. Berletch, J. B. et al. Escape from X Inactivation Varies in Mouse Tissues. PLoS Genet. 11, (2015).

52. Tukiainen, T. et al. Landscape of X chromosome inactivation across human tissues. Nature 550, 244– 248 (2017).

53. Grubman, A. et al. A single-cell atlas of entorhinal cortex from individuals with Alzheimer’s disease reveals cell-type-specific gene expression regulation. Nat. Neurosci. 22, 2087–2097 (2019).

54. Morabito, S. et al. Single-nucleus chromatin accessibility and transcriptomic characterization of Alzheimer’s disease. Nat. Genet. 53, 1143–1155 (2021).

55. Sternson, S. M. Hypothalamic Survival Circuits: Blueprints for Purposive Behaviors. Neuron 77, 810–824 (2013).

56. Leibowitz, S. F. & Wortley, K. E. Hypothalamic control of energy balance: different peptides, different functions. Peptides 25, 473–504 (2004).

57. Kleshchevnikov, V. et al. Cell2location maps fine-grained cell types in spatial transcriptomics. Nat. Biotechnol. 1–11 (2022) doi:10.1038/s41587-021-01139-4.

58. Fricker, L. D. et al. Identification and Characterization of proSAAS, a Granin-Like Neuroendocrine Peptide Precursor that Inhibits Prohormone Processing. J. Neurosci. 20, 639–648 (2000).

59. Cao, J. et al. The single-cell transcriptional landscape of mammalian organogenesis. Nature 566, 496–502 (2019).

60. Vann, S. D. & Nelson, A. J. D. The mammillary bodies and memory. in Progress in Brain Research vol. 219 163–185 (Elsevier, 2015).

61. Chrousos, G. P. Regulation and Dysregulation of the Hypothalamic-Pituitary-Adrenal Axis: The Corticotropin-Releasing Hormone Perspective. Endocrinol. Metab. Clin. North Am. 21, 833–858 (1992).

62. Mouradian, M. M. et al. Spinal fluid CRF reduction in Alzheimer’s disease. Neuropeptides 8, 393– 400 (1986).

63. Bayatti, N. & Behl, C. The neuroprotective actions of corticotropin releasing hormone. Ageing Res. Rev. 4, 258–270 (2005).

64. Chen, T. & He, T. Xgboost: extreme gradient boosting. (2015).

65. Lundberg, S. & Lee, S.-I. A Unified Approach to Interpreting Model Predictions. ArXiv170507874 Cs Stat (2017).

66. Leng, K. et al. Molecular characterization of selectively vulnerable neurons in Alzheimer’s disease. Nat. Neurosci. 24, 276–287 (2021).

67. Burger, J. M. S. Sex-Specific Effects of Interventions That Extend Fly Life Span. Sci. Aging Knowl. Environ. 28, 30 (2004).

68. Honjoh, S., Ihara, A., Kajiwara, Y., Yamamoto, T. & Nishida, E. The Sexual Dimorphism of Dietary Restriction Responsiveness in Caenorhabditis elegans. Cell Rep. 21, 3646–3652 (2017).

69. Davis, E. J., Lobach, I. & Dubal, D. B. Female XX sex chromosomes increase survival and extend lifespan in aging mice. Aging Cell 18, e12871 (2019).

70. Creamer, K. M., Kolpa, H. J. & Lawrence, J. B. Nascent RNA scaffolds contribute to chromosome territory architecture and counter chromatin compaction. Mol. Cell 81, 3509–3525.e5 (2021).

71. Stuart, T. et al. Comprehensive Integration of Single-Cell Data. Cell 177, 1888–1902.e21 (2019).

72. Liberzon, A. et al. The Molecular Signatures Database (MSigDB) hallmark gene set collection. Cell Syst. 1, 417–425 (2015).

73. Kuleshov, M. V. et al. Enrichr: a comprehensive gene set enrichment analysis web server 2016 update. Nucleic Acids Res. 44, W90–W97 (2016).

74. Benarroch, E. E. Oxytocin and vasopressin: Social neuropeptides with complex neuromodulatory functions. Neurology 80, 1521–1528 (2013).

75. Veenema, A. H. & Neumann, I. D. Central vasopressin and oxytocin release: regulation of complex social behaviours. in Progress in Brain Research (eds. Neumann, I. D. & Landgraf, R.) vol. 170 261–276 (Elsevier, 2008).

76. Wittmann, W. et al. Prodynorphin-Derived Peptides Are Critical Modulators of Anxiety and Regulate Neurochemistry and Corticosterone. Neuropsychopharmacology 34, 775–785 (2009).

77. Mclaughlin, C., Baile, C., Dellafera, M. & Kasser, T. Meal-stimulated increased concentrations of CCK in the hypothalamus of Zucker obese and lean rats☆. Physiol. Behav. 35, 215–220 (1985).

78. Hiller-Sturmhöfel, S. & Bartke, A. The Endocrine System. Alcohol Health Res. World 22, 153–164 (1998).

79. Nemeroff, C. B. et al. Neurotensin: Central nervous system effects of a hypothalamic peptide. Brain Res. 128, 485–496 (1977).

80. Sapouckey, S. A., Deng, G., Sigmund, C. D. & Grobe, J. L. Potential mechanisms of hypothalamic renin-angiotensin system activation by leptin and DOCA-salt for the control of resting metabolism. Physiol. Genomics 49, 722–732 (2017).

81. Gardiner, J. V. et al. Prokineticin 2 Is a Hypothalamic Neuropeptide That Potently Inhibits Food Intake. Diabetes 59, 397–406 (2010).

82. Navarro, V. M. et al. The Integrated Hypothalamic Tachykinin-Kisspeptin System as a Central Coordinator for Reproduction. Endocrinology 156, 627–637 (2015).

83. Braasch, D. C., Deegan, E. M., Grimm, E. R. & Griffin, J. D. Calcitonin gene-related peptide alters the firing rates of hypothalamic temperature sensitive and insensitive neurons. BMC Neurosci. 9, 64 (2008).

84. Osterstock, G. et al. Somatostatin triggers rhythmic electrical firing in hypothalamic GHRH neurons. Sci. Rep. 6, 24394 (2016).

85. Taylor, T., Wondisford, F. E., Blaine, T. & Weintraub, B. D. The paraventricular nucleus of the hypothalamus has a major role in thyroid hormone feedback regulation of thyrotropin synthesis and secretion. Endocrinology 126, 317–324 (1990).

86. De Lecea, L. & Huerta, R. Hypocretin (orexin) regulation of sleep-to-wake transitions. Front. Pharmacol. 5, 16 (2014).

87. Sakurai, T. et al. Orexins and orexin receptors: a family of hypothalamic neuropeptides and G protein-coupled receptors that regulate feeding behavior. Cell 92, 573–585 (1998).

88. Madelaine, R. et al. The hypothalamic NPVF circuit modulates ventral raphe activity during nociception. Sci. Rep. 7, 41528 (2017).

89. Chartrel, N. et al. The Neuropeptide 26RFa (QRFP) and Its Role in the Regulation of Energy Homeostasis: A Mini-Review. Front. Neurosci. 10, (2016).

90. Chen, A. et al. QRFP and Its Receptors Regulate Locomotor Activity and Sleep in Zebrafish. J. Neurosci. 36, 1823–1840 (2016).

91. Sakata, T., Yoshimatsu, H. & Kurokawa, M. Hypothalamic neuronal histamine: Implications of its homeostatic control of energy metabolism. Nutrition 13, 403–411 (1997).

92. Kiaris, H., Chatzistamou, I., Papavassiliou, A. G. & Schally, A. V. Growth hormone-releasing hormone: not only a neurohormone. Trends Endocrinol. Metab. 22, 311–317 (2011).

93. Gamble, K. L., Allen, G. C., Zhou, T. & McMahon, D. G. Gastrin-Releasing Peptide Mediates Light-Like Resetting of the Suprachiasmatic Nucleus Circadian Pacemaker through cAMP Response Element-Binding Protein and Per1 Activation. J. Neurosci. 27, 12078–12087 (2007).

94. Romanov, R. A. et al. Molecular interrogation of hypothalamic organization reveals distinct dopamine neuronal subtypes. Nat. Neurosci. 20, 176–188 (2017).

95. Vosko, A. M., Schroeder, A., Loh, D. H. & Colwell, C. S. Vasoactive intestinal peptide and the mammalian circadian system. Gen. Comp. Endocrinol. 152, 165–175 (2007).

